# Proliferation history and transcription factor levels drive direct conversion

**DOI:** 10.1101/2023.11.26.568736

**Authors:** Nathan B. Wang, Brittany A. Lende-Dorn, Honour O. Adewumi, Adam M. Beitz, Patrick Han, Timothy M. O’Shea, Kate E. Galloway

## Abstract

The sparse and stochastic nature of reprogramming has obscured our understanding of how transcription factors drive cells to new identities. To overcome this limit, we developed a compact, portable reprogramming system that increases direct conversion of fibroblasts to motor neurons by two orders of magnitude. We show that subpopulations with different reprogramming potentials are distinguishable by proliferation history. By controlling for proliferation history and titrating each transcription factor, we find that conversion correlates with levels of the pioneer transcription factor Ngn2, whereas conversion shows a biphasic response to Lhx3. Increasing the proliferation rate of adult human fibroblasts generates morphologically mature, induced motor neurons at high rates. Using compact, optimized, polycistronic cassettes, we generate motor neurons that graft with the murine central nervous system, demonstrating the potential for *in vivo* therapies.

**One Sentence Summary:** Using a systems and synthetic biology approach to study the molecular determinants of reprogramming, we find that proliferation history and transcription factor levels drive cell fate in direct conversion to motor neurons.

**Highlights:**

- minimal high-efficiency cocktail allows systematic interrogation of the reprogramming process
- history provides a principal axis to distinguish transcription factors’ influence
- of the transcription factor cocktail impacts reprogramming efficiency and dynamics
- of individual transcription factors differentially influence the rate of reprogramming
- early hyperproliferation increases direct conversion of adult human fibroblasts
- ptimal cocktail allows neurotrophic factor-free reprogramming and *in vivo* grafting

## Introduction

The inherently stochastic nature of reprogramming has limited the identification of design rules for guiding cells through cell-fate transitions (Shakiba et al., 2019; Yamanaka, 2009). Combinations of transcription factors guide cells to specific new cell fates (Chen et al., 2021; Feng et al., 2008; Ieda et al., 2010; Lujan et al., 2012; Mall et al., 2017; Rosa et al., 2018; Son et al., 2011; Soufi et al., 2015, 2012; Sridharan et al., 2009; Takahashi and Yamanaka, 2006; Velychko et al., 2019; Wang Li et al., 2015; Wernig et al., 2008). However, within these diverse cocktails, how the individual transcription factors and their stoichiometries influence reprogramming events remains poorly defined (An et al., 2019; Carey et al., 2011; Ieda et al., 2010; Kempf et al., 2021; Velychko et al., 2019, 2019; Wang Li et al., 2015).

Extrinsic variation including variability across batches of cells, complex cocktails of transcription factors, and expression of those factors, as well as intrinsic cellular variation have obscured how the levels of individual transcription factors promote or impede reprogramming (Jain et al., 2023; Wang et al., 2020). Process variability combined with low rates of reprogramming constrain our ability to resolve the distinct molecular states and processes that cells adopt during successful reprogramming (Ilia et al., 2023). By improving the rate of reprogramming, we can identify these states and processes that promote cell-fate transitions.

Across diverse systems, previous work has established that higher expression of reprogramming factors correlates with higher rates of reprogramming (Babos et al., 2019; Ilia et al., 2023; Jindal et al., 2023). However, inclusion or dropout of individual transcription factors influences the number of reprogramming events, transcriptional trajectories, and function of reprogrammed cells (Hersbach et al., 2022; Mall et al., 2017; Velychko et al., 2019). While transcriptional profiling identifies expression trajectories, transcription factors exert their influence as proteins, which cannot be directly imputed from mRNA expression (Hafner et al., 2017; Ideker et al., 2001; Lundberg et al., 2010). Moreover, global processes such as transcription, replication, and proliferation strongly influence reprogramming events and may amplify or dampen the expression and activity of transcription factors (Babos et al., 2019; Hu et al., 2020; Jain et al., 2023; Kueh et al., 2013; Palozola et al., 2017; Percharde et al., 2017; Stadhouders et al., 2018). Thus, transcription factors are a powerful driver of cellular reprogramming but how they combine with other global processes to guide cell-fate transitions remains unclear.

During development, progenitor cells often divide prior to differentiation (Bowman et al., 2008; Bultje et al., 2009; Eriksson et al., 1998). Division supports differentiation and maintenance of the progenitor pool as cells adopt distinct fates. In direct conversion, cells do not necessarily transit through a defined progenitor state (Son et al., 2011) and proliferation is not required for reprogramming (Di Tullio and Graf, 2012; Fishman et al., 2015; Heinrich et al., 2010). However, highly proliferative cells reprogram at higher rates to both induced pluripotent stem cells (iPSCs) and to post-mitotic neurons, suggesting that proliferation may serve a conserved role in facilitating cell-fate transitions (Babos et al., 2019; Guo et al., 2014; Jain et al., 2023). Curiously, we and others have observed that the rate of proliferation can substantially impact the expression of transcription factors in both native and synthetic systems (Babos et al., 2019; Kueh et al., 2013). Indeed, in differentiation to macrophages, cell-cycle lengthening enables progenitors to commit to the macrophage lineage by increasing accumulation of PU.1 protein without increasing PU.1 transcription (Kueh et al., 2013). If higher rates of proliferation simultaneously increase the probability of reprogramming and reduce expression of the transcription factors, how do cells integrate these incoherent cues?

To examine how individual transcription factors and their levels of expression contribute to reprogramming outcomes, we selected a model of direct conversion from fibroblasts to motor neurons (Babos et al., 2019). Direct conversion to motor neurons provides a well-defined testbed with established murine transgenic reporters for staging the reprogramming process in living cells. Importantly, direct conversion to a post-mitotic identity provides several advantages over reprogramming to iPSCs. Conversion to a post-mitotic cell type such as motor neurons allows us to decouple transient changes in proliferation that facilitate cell-fate transitions from proliferation associated with the new cell identity. Further, we can accurately estimate reprogramming events based simply on the number of neurons generated which is not possible in reprogramming to iPSCs. Unlike iPSCs, neurons do not divide once reprogrammed. Thus, each neuron corresponds to exactly one reprogramming event.

While post-mitotic neurons are useful for quantifying reprogramming events, direct conversion protocols to non-proliferative cell fates often suffer from low reprogramming yields. To better dissect how transcription factor levels direct cell-fate transitions, we addressed the challenge of low reprogramming rates by capitalizing on our recently developed chemo-genetic cocktail that increases direct conversion to motor neurons by two orders of magnitude (Babos et al., 2019). Importantly, we first sought to reduce sources of extrinsic variation to resolve how individual transcription factors influence reprogramming outcomes. To this end, we tailored the transcription factor cocktail and minimized the number of viruses, consequently increasing the fraction of cells that undergo a period of rapid proliferation. These cells with a hyperproliferative history reprogram at five-fold higher rates than non-hyperproliferative cells, leading to an exponential increase in yield. By increasing the number of reprogramming events and titrating expression, we identified transcription factor-specific correlations between expression and reprogramming events. Using these insights, we designed improved reprogramming cocktails for direct conversion of adult human fibroblasts to motor neurons. To demonstrate translational potential, we developed minimal vector systems that generate motor neuron grafts capable of integrating with the murine central nervous system. Overall, our results show that proliferation history and transcription factor expression combine to drive cell-fate transitions.

## Results

### Tailored conversion cocktail minimizes extrinsic variation and increases reprogramming events

We aimed to minimize sources of extrinsic variation so we could precisely dissect the processes that support cell-fate transitions. Capitalizing on our recently developed high-efficiency cocktail (DDRR) (Babos et al., 2019), we used the conversion of mouse embryonic fibroblasts (MEFs) to induced motor neurons (iMNs) as our model system. We delivered the reprogramming factors encoded on retroviruses to transgenic MEFs bearing the motor neuron reporter, Hb9::GFP (Wichterle et al., 2002), which allows us to monitor reprogramming dynamically. Activation of Hb9::GFP and neuronal morphology provide live metrics of the conversion process (Fig. 1A-B, S1A). Importantly, each fully reprogrammed induced motor neuron corresponds to exactly one reprogramming event, allowing us to quantitatively evaluate how perturbations impact the number of successful events.

**Fig. 1.**
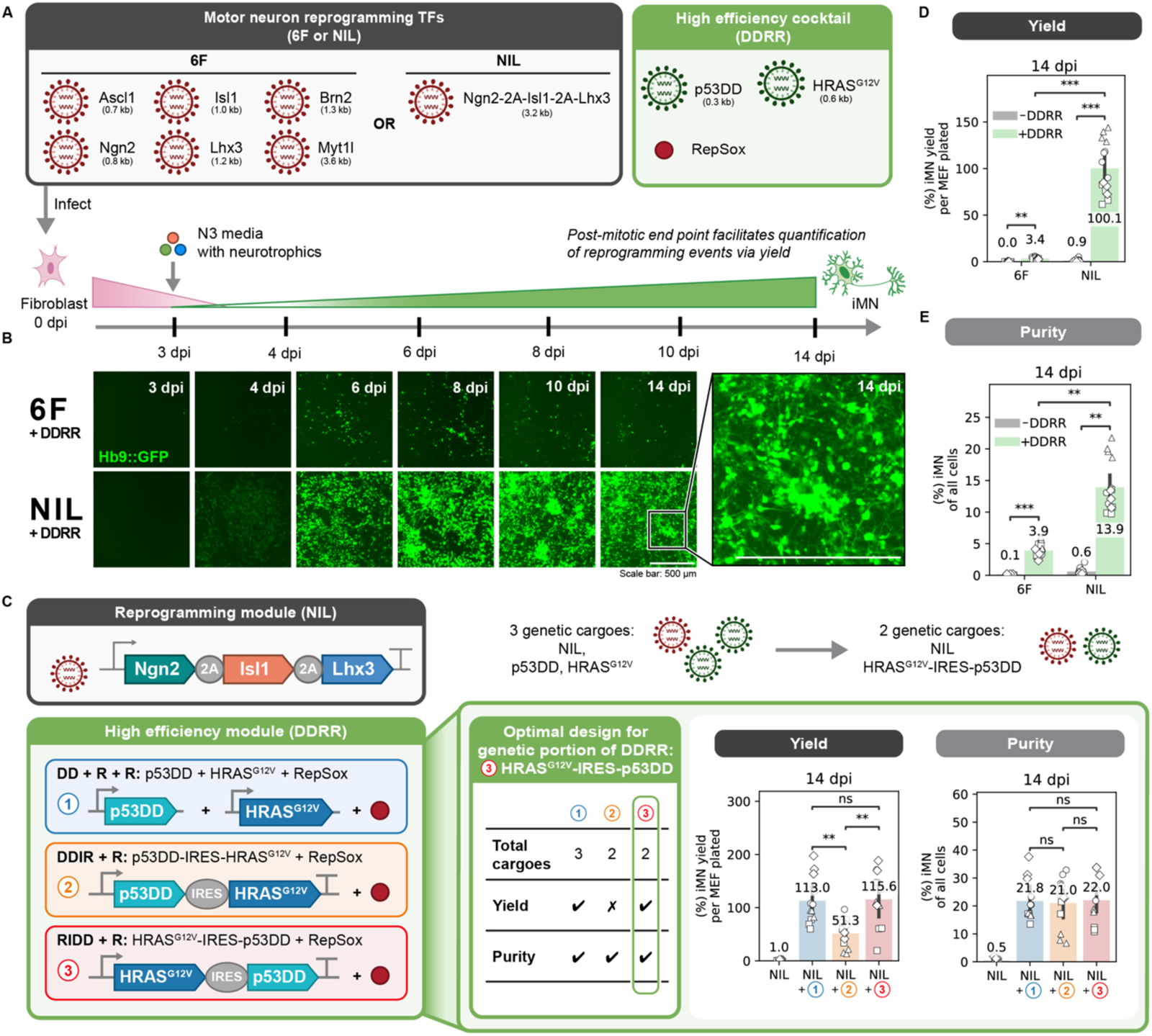
Tailored conversion cocktail minimizes extrinsic variation and increases reprogramming events. A. Schematic depicting the reprogramming process. B. Images of Hb9::GFP activation and morphological changes of MEF-to-iMN reprogramming starting at 3 days post infection (dpi) until 14 dpi when reprogramming purity and yield are quantified. Inset shows zoomed in portion of image. Both scale bars represent 500 µm. C. Reprogramming purity (left) and yield (right) at 14 dpi for different designs of the genetic portion of the DDRR cocktail. Mean is shown with 95% confidence interval; marker styles denotes bio reps; n = 4 biological reps per condition; two-tailed t-test with Bonferroni correction. D-E. Reprogramming purity (D) and yield (E) at 14 dpi for 6F vs. NIL ± DDRR. Mean is shown with 95% confidence interval; marker styles denotes biological reps; n = 4 biological reps per condition; one-tailed t-test with Bonferroni correction. Significance summary: p > 0.05 (ns); *p ≤ 0.05; **p ≤ 0.01; ***p ≤ 0.001; and ****p ≤ 0.0001

While our previously published protocol improved rates of direct conversion, we observed substantial variability across biological replicates (Babos et al., 2019). We suspected the variability in reprogramming rate was due to extrinsic variation introduced by cotransduction of eight different viruses. To reduce extrinsic variation and improve the controllability, we minimized the number of transcription factors and viruses. First, we sought to identify the minimal set of transcription factors from the originally identified six factors (6F: Ascl1, Brn2, Myt1l, Ngn2, Isl1, and Lhx3) with our previously identified high-efficiency cocktail (Babos et al., 2019; Son et al., 2011). Induction of Ngn2, Isl1, and Lhx3 expression efficiently drives mouse embryonic stem cells to motor neurons (Mazzoni et al., 2013). Further, Ngn2, Isl1, and Lhx3 (NIL) can be expressed from a single-transcript cassette where each transcription factor is separated by “self-cleaving” 2A sequences (Mazzoni et al., 2013). As a 3.2 kilobase cargo, these three transcription factors can be efficiently delivered by a single retrovirus to induce reprogramming (Fig. 1A). We verified that Ngn2, Isl1, and Lhx3 together comprise a minimally sufficient set for high-efficiency MEF-to-iMN direct conversion through a transcription factor dropout experiment (Fig. S1B-C). Surprisingly, NIL accelerated the number of Hb9::GFP positive cells that formed beginning at 4 days post-infection (4 dpi) compared to 6F with the high-efficiency DDRR cocktail (Fig. 1B).

Next, we encoded the transgenes in the high-efficiency DDRR module into a single viral cargo. We explored how the placement of p53DD and HRAS^G12V^ relative to an internal ribosome entry site (IRES) impacts the rate of conversion. To capture multiple dimensions of the reprogramming process, we quantified reprogramming rates using two metrics: yield and purity. We define yield as how many reprogrammed cells are at the final time point relative to the starting cell input. Yield increases for highly proliferative conditions and measures how many successful reprogramming events occur from an initial seeded population. We also define purity as the percent of cells of the total population that is reprogrammed at that time point. Purity measures the proportion of the cells that have converted into the desired cell fate. Overexpressing the DDRR module variants with the NIL reprogramming module, we observed similar reprogramming purities. However, the yield doubles for the RIDD construct where HRAS^G12V^ is placed upstream, compared to when the DDIR construct where p53DD is upstream (Fig. 1C).

With our optimal DDRR design (RIDD), we returned to our reprogramming transcription factor module and compared the 6F cocktail with the single polycistronic cargo, NIL. Compared to 6F, NIL increases the reprogramming yield 30-fold and purity 4-fold (Fig. 1D-E). Having identified a highly efficient and minimal set of genetic cargoes, we used the optimal RIDD design for the high-efficiency DDRR module to systematically examine how individual transcription factors impact reprogramming. We began with examining the basis for higher reprogramming efficiency with NIL over 6F. By reducing extrinsic sources of variability, we developed a highly robust process for the direct conversion of fibroblasts to post-mitotic motor neurons, providing us a tool to systematically interrogate how transcription factor levels correlate to reprogramming events.

### Proliferation history provides a principal axis to distinguish transcription factors’ influence

Hyperproliferative cells reprogram at higher rates to both mitotic and post-mitotic identities, establishing hyperproliferation as a powerful driver of diverse cell-fate transitions (Babos et al., 2019; Guo et al., 2014; Jain et al., 2023). Potentially, different conversion rates induced by different transcription factor cocktails can be attributed to cells integrating a differential set of molecular interactions that lead to different rates of proliferation.

To examine whether differences in proliferation rates might distinguish cocktails’ reprogramming rates, we measured the proliferation history of cells undergoing reprogramming in the presence of different reprogramming cocktails. Cells that undergo a period of hyperproliferation were identified by assaying dilution of CellTrace dye for 72 hours from 1 to 4 dpi, an early timepoint in iMN direct conversion, as previously established (Babos et al., 2019) (Fig. 2A). We denote cells with a history of rapid proliferation at 4 dpi as hyperproliferative (hyperP) and all other cells at 4 dpi as non-hyperP. We quantify hyperproliferation by both the percent of cells in the population that are hyperproliferative and the total number of hyperproliferative cells. As expected, the fraction and total number of hyperproliferative cells scale exponentially (Fig. 2B, S2H). In addition, as we expect from its higher reprogramming efficiency, NIL + DDRR has a higher fraction and number of hyperproliferative cells than 6F + DDRR (Fig. S2A-B).

**Fig. 2.**
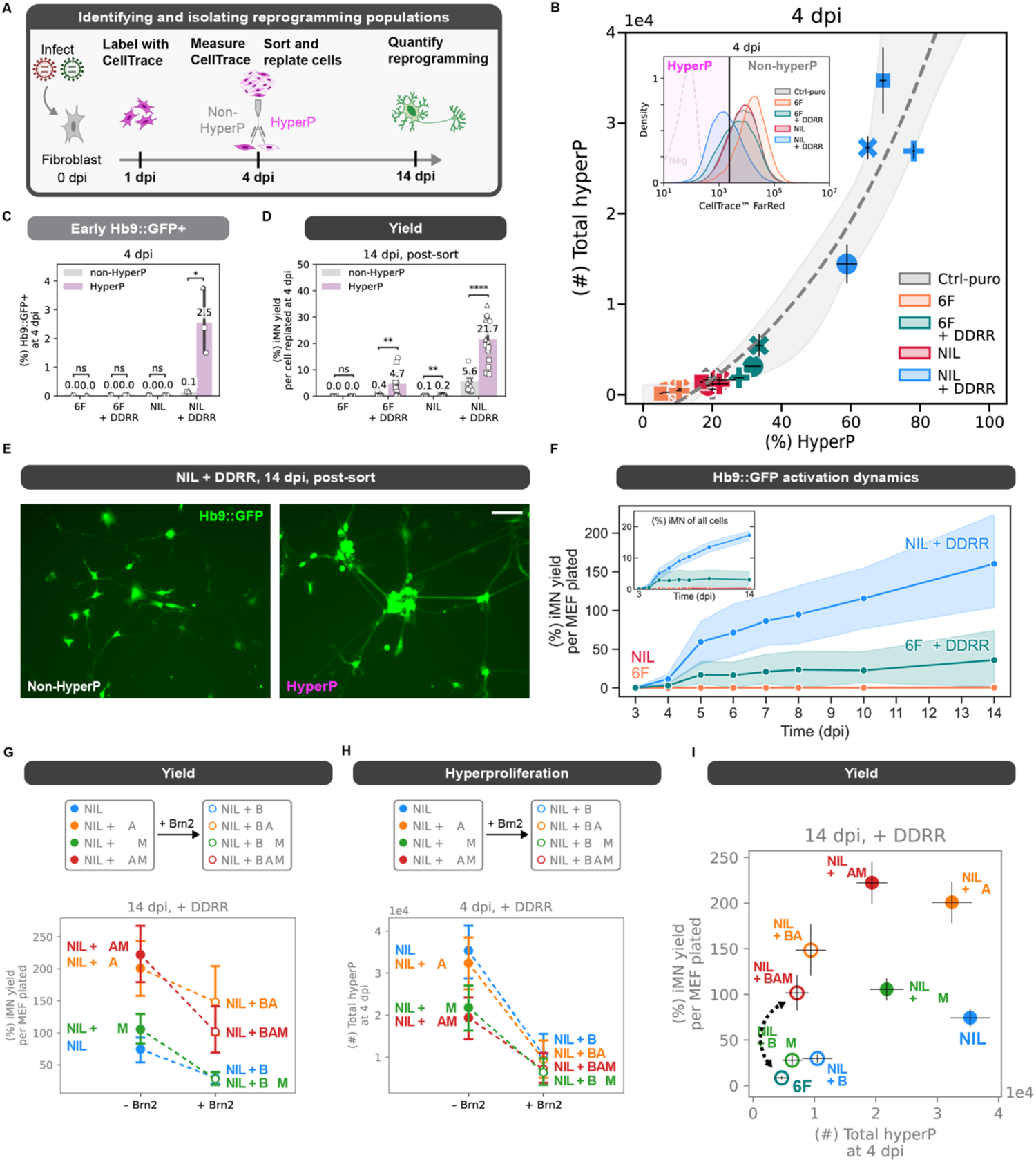
Proliferation history provides a principal axis to distinguish transcription factors’ influence. A. Schematic of the CellTrace dilution assay. Cells are labeled at 1 dpi and their fluorescent intensity is measured at 4 dpi, where cells that have gone through many divisions have low fluorescence. For sorting experiments, cells are sorted at 4 dpi into hyperP and non-hyperP according to their fluorescent intensity relative to a control puro infected condition for a given biological replicate. Cells are replated on 4 dpi at the same initial seeding density at 10k cells/96-well. B. HyperP total number vs. percent at 4 dpi across all reprogramming conditions. An exponential fit is shown with 95% confidence interval. Representative CellTrace distributions across reprogramming conditions. Hyperproliferative cells are defined relative to the 20%-lowest fluorescent cells in a control puro (Ctrl-puro) infected condition for a given biological replicate. C. Early Hb9::GFP reporter expression at 4 dpi in hyperP vs. non-hyperP populations within each condition. Mean is shown with 95% confidence interval; marker styles denotes biological reps; n = 3 biological reps per condition; one-tailed t-test. D. Reprogramming yield at 14 dpi for cells sorted and replated at the same initial seeding density at 10k cells/96-well into hyperP vs. non-hyperP populations within each condition. Mean is shown with 95% confidence interval; marker styles denotes biological reps; n = 3 biological reps per condition; one-tailed t-test. E. Representative images at 14 dpi of NIL + DDRR reprogrammed cells that were sorted from hyperP and non-hyperP at 4 dpi. Scalebar represents 100 µm. F. Yield of cells activating Hb9::GFP reporter expression from 3-14 dpi for 6F vs. NIL ± DDRR. Inset shows the purity over time. Mean is shown with 95% confidence interval. n = 3 biological reps per condition. G-H. Effect of adding in Brn2 on reprograming yield (G) and hyperproliferation at 4 dpi (H) for each reprogramming cocktail (all reprogrammed with +DDRR). Dashed lines connect cocktails without Brn2 (closed circle) to corresponding ones with Brn2 (open circle). Mean is shown ± 95% CI; n = 4 biological reps per condition. I. Reprogramming purity at 14 dpi vs. toal number of hyperP cells at 4 dpi for each reprogramming cocktail (all reprogrammed with +DDRR). Mean is shown ± standard error of mean (SEM); n = 4 biological reps per condition. Black dashed line connects 6F when NIL is encoded on 3 separate viruses (i.e. A+B+M+N+I+L) vs. when NIL is on a single polycistronic transcript (i.e. A+B+M+NIL). Significance summary: p > 0.05 (ns); *p ≤ 0.05; **p ≤ 0.01; ***p ≤ 0.001; and ****p ≤ 0.0001

To examine the rates of conversion for cells with different proliferation histories, we isolated and replated the hyperproliferative and non-hyperproliferative cells at 4 dpi, and measured conversion at 14 dpi (Fig. 2A, S2C-D). Compared to non-hyperproliferative cells, hyperproliferative cells reprogrammed with NIL + DDRR were 25-fold more likely to express Hb9::GFP early on at 4 dpi (Fig. 2C), and achieve higher reprogramming yield and purity (Fig. 2D, Fig. S2E). By normalizing yield to the number of cells replated at 4 dpi, we can observe the propensity of cells to convert based on proliferation history. A history of early hyperproliferation increases the probability of cells successfully converting by 4-fold in NIL+ DDRR. Further, compared to non-hyperproliferative cells, hyperproliferative cells adopted a more mature, neuronal morphology at 14 dpi, displaying compact, regular soma with interconnected networks of axons (Fig. 2E, Fig. S2D). Taken together, these data show that hyperproliferation not only generates more cells, but also the cells generated during hyperproliferation possess a greater propensity to reprogram and develop mature morphologies.

We hypothesized that specific DDRR cassette designs may increase the rate of proliferation and thus the rates of early Hb9::GFP activation and of reprogramming (Fig. 1C). As expected, increased numbers of hyperproliferative cells yields higher numbers of Hb9::GFP cells at 4 and 14 dpi (Fig. S2F-G). Curiously, the fraction of Hb9::GFP positive cells at 14 dpi remains unchanged across DDRR variants (Fig. S2G-H). Putatively, the encoding of p53DD and HRAS^G12V^ generates different reprogramming rates due to their effects on proliferation.

We wondered whether increased rates of hyperproliferation in NIL compared to 6F coincide with a more rapid adoption of the iMN cell fate. To assess the dynamics of reprogramming events, we tracked Hb9::GFP expression over the reprogramming process. When DDRR is included with either the NIL or 6F cocktails, Hb9::GFP activation begins at 4 dpi (Fig. 1B). For 6F + DDRR, the number and fraction of Hb9::GFP positive cells plateaus around 5 dpi. Thus, most conversion events occur early with 6F + DDRR (Fig. 2F). In contrast, cells reprogrammed with NIL + DDRR continue to generate Hb9::GFP positive cells throughout the reprogramming process. We confirmed that adding DDRR to the base NIL cocktail induces a transient window of hyperproliferation by immunofluorescent staining of Ki67, a marker of proliferation (Fig. S2I-L). By 14 dpi, with or without DDRR, the majority of Hb9::GFP positive cells do not express Ki67, indicating they have exited cell cycle and adopted a post-mitotic identity (Fig. S2K,M-N). Expanding the conversion window not only increases the number of cells but increases the fraction of cells that fully reprogram (Fig 2F). Altogether, our data suggests a model in which driving proliferation provides an exponential benefit in the conversion to induced motor neurons. Proliferation not only expands the number of cells but also endows this expanded population with a greater propensity to reprogram.

### Transcription factor identity and encoding impact conversion rates and dynamics

As transcription factors can work synergistically, we wondered if differences in reprogramming rates between the 6F and NIL cocktails were due to a single or multiple transcription factors. Given that the 6F cocktail reduces proliferation more than NIL in the presence of the high-efficiency DDRR module (Fig. S2A-B), we hypothesized that one or multiple of the three BAM (Brn2, Asl1, Myt1L) transcription factors may impede proliferation. Using the polycistronic NIL with the high-efficiency DDRR cocktail as the reference, we added in all six possible combinations of the three BAM factors to assess their individual and combined effects on proliferation and reprogramming. Inclusion of Brn2 reduces proliferation, activation of Hb9::GFP at 4 dpi, and conversion yield and purity (Fig. 2G-I, Fig. S2O-Q). In contrast, addition of Ascl1 and Myt1l increases iMN yield without altering proliferation (Fig. 2I, S2P). Together, these data suggest that individual transcription factors can independently or synergistically influence proliferation and reprogramming.

Notably, the encoding of the TF cocktail, not just the identity of the constituents, impacts reprogramming. The cocktails of 6F and NIL + BAM contain an identical set of transcription factors. However, in 6F, Ngn2, Isl1, and Lhx3 are expressed from separate viruses whereas NIL expresses the same factors from a single transcript. The NIL + BAM cocktail reprograms with 7-fold higher purity and 12-fold higher yield while minimally impacting proliferation (Fig. 2I). These data suggest that transcription factor encoding can substantially improve (or hinder) cocktail efficacy. Putatively, the single transcript design may improve efficacy by controlling for co-delivery and stoichiometry and/or by reducing the number of viruses. Together, these observations demonstrate that while the identity of the transcription factors strongly influences reprogramming, the encoding and delivery of those factors impact the frequency and likelihood of a reprogramming event. With that in mind, we sought to determine how each of the three motor neuron transcription factors (Ngn2, Isl1, and Lhx3) influences the number of induced motor neurons.

### Titration of individual transcription factors reveals factor-specific influence on conversion rates

To investigate how the levels of ectopic transcription factor expression influence reprogramming, we titrated individual transcription factors, keeping the other genetic cargos constant (Fig. 3A-B). We used arrays of small upstream open reading frames (uORFs) to generate a range of individual transcription factor levels while keeping the retroviral delivery vehicle the same. uORFs change gene expression by altering translation rates (Ferreira et al., 2013; Jones et al., 2020). By adding a fluorescent 2A-mRuby2 tag downstream of the transcription factor (Fig. 3B), we built single-cell, live reporters of ectopic transcription factor levels. The mRuby2 levels provide a proxy for the ectopic expression of the linked transcription factor early in the reprogramming process at 4 dpi. Using these tools, we titrated the level of each transcription factor while keeping other factors and the high-efficiency DDRR module constant.

**Fig. 3.**
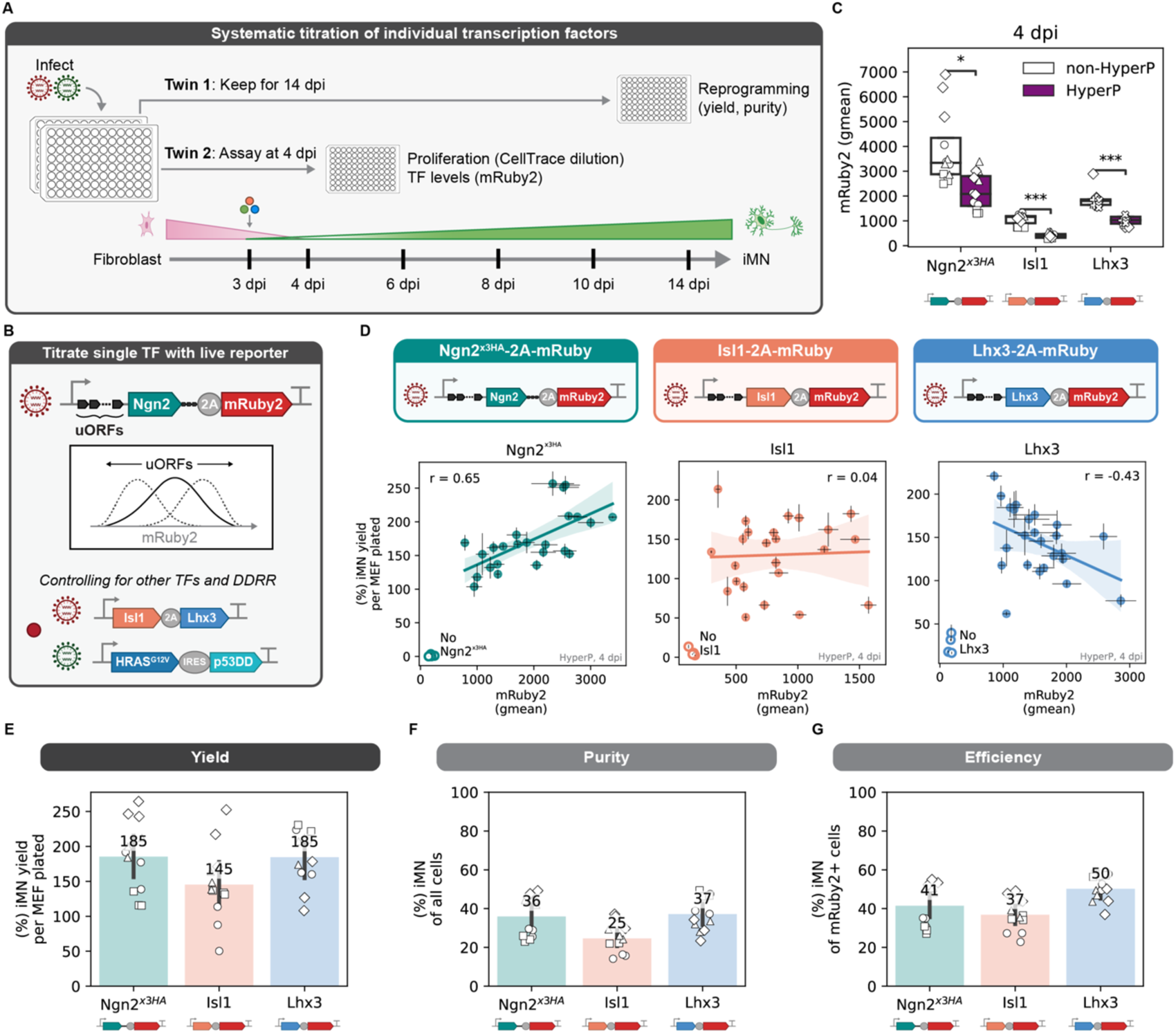
Titration of individual transcription factors reveals factor-specific influence on conversion rates. A. Overview of coupled time-point (4 and 14 dpi) approach for titrating transcription factor expression. Cells are reprogrammed in biological duplicates where one twin is assayed at 4 dpi to measure expression of the mRuby2 reporter in hyperproliferative cell. The other twin is kept until 14 dpi to measure reprogramming outcomes using Hb9::GFP expression measured via flow cytometry. B. Schematic of how individual transcription factor levels were titrated and measured using upstream open reading frames (uORFs) placed upstream of the reprogramming transcription factor of interest with a 2A-mRuby2 fluorescent reporter. All other factors were kept constant, *e.g.* if Ngn^x3HA^-2A-mRuby2 is titrated, then Isl1, Lhx3, and +DDRR expression are not altered using uORFs. C. Expression of transcription factor of interest with a 2A-mRuby2 fluorescent reporter in non-hyperproliferative (non-HyperP) and hyperproliferative (HyperP) cells at 4 dpi. Cells are reprogrammed with non-fluorescent versions of the other transcription factors (*e.g.* Ngn^x3HA^-2A-mRuby2 is reprogrammed with Isl1, Lhx3, and +DDRR, see Fig. 3B). Geometric mean is shown by markers with a box plot; marker styles denote biological reps; n = 4 biological reps per condition; one-tailed t-test. D. iMN yield at 14 dpi vs. mRuby2 expression in just the hyperP cells at 4 dpi to avoid confounding proliferation with expression across all uORFs for Ngn^x3HA^-2A-mRuby2 (left), Isl1-2A-mRuby2 (middle), and Lhx3-2A-mRuby2 (right). Open marker denotes the conditions where the transcription factor-2A-mRuby2 being titrated is excluded but all other factors are included. Mean is shown ± s.e.m. for each uORF for a single biological rep; n = 4 biological reps per condition. E-G. iMN yield (E), purity (F), and efficiency (G) at 14 dpi for each transcription factor-2A-mRuby2 fluorescent reporter with no uORFs. Cells are reprogrammed with non-fluorescent versions of the other transcription factors (*e.g.* Ngn^x3HA^-2A-mRuby2 is reprogrammed with Isl1, Lhx3, and +DDRR, see Fig. 3B). Marker styles denote biological reps; n = 4 biological reps per condition. Significance summary: p > 0.05 (ns); *p ≤ 0.05; **p ≤ 0.01; ***p ≤ 0.001; and ****p ≤ 0.0001

Consistent with our previous observations, the expression of each of the three transcription factor-2A-mRuby2 transgenes are lower in cells that undergo a period of hyperproliferation compared to cells that do not (Fig. 3C). Because we found hyperproliferative cells are 25-fold more likely to activate Hb9::GFP early and 4-fold more likely to convert to morphologically mature, motor neurons (Fig. 2C-E), we chose to examine the transgene expression within this population of hyperproliferative cells. This allowed us to eliminate the confounder of different levels of expression across subpopulations of cells. Thus, we hypothesized that trends observed in these cells may more clearly reveal the influence of transcription factor levels on reprogramming events.

To obtain a range of expression levels, we introduced five different uORFs upstream of the coding sequence of each transcription factor encoded with a mRuby2 reporter. To exploit natural biological heterogeneity in expression and control for variance across reprogramming biological replicates, we paired reprogramming assays to generate a variety of expression profiles at 4 dpi with a corresponding biological twin to measure reprogramming outcomes at 14 dpi (Fig. 3A). Importantly, systematically changing transcription factor expression in our tailored reprogramming module does not affect proliferation rates (Fig. S3A). Thus, our designs remove differences in proliferation history as a confounding variable across the genetic variants. As expected, introducing uORFs to transgenes preserves the trend of lower expression in the hyperproliferative cells (Fig. S3B).

Utilizing our paired reprogramming assays, we find that high levels of ectopic Ngn2^x3HA^ leads to improved reprogramming yields and ectopic Isl1 is uncorrelated (Fig. 3D, S3C). Comparing all three transcription factor reporters as controls for each other, reprogramming potential does not simply correlate with high expression of the mRuby2 reporter, suggesting the transcription factor identity sets the relationship between expression and reprograming. The positive correlation of reprogramming yield with expression of the pioneer factor, Ngn2^x3HA^, mirrors observations from iPSC reprogramming where high expression of the pioneer factor, Oct4, increases the rate of colony formation (Velychko et al., 2019). As expected, Ngn2^x3HA^-2A-mRuby2 expression varies across biological replicates, yet within each biological replicate, the correlation between yield and levels of Ngn2 is preserved (Fig. S3C). Surprisingly, we find that ectopic Lhx3 induces a biphasic response in reprogramming yield (Fig. 3D). In the absence of ectopic Lhx3, conversion rates are low. However, high expression of ectopic Lhx3 decreases reprogramming yield, demonstrating a biphasic reprogramming response to Lhx3. Putatively, moderate levels of Lhx3 expression support optimal reprogramming. Together, our data suggest that in the presence of Isl1, a combination of high levels of Ngn2 and low levels of Lhx3 promote reprogramming to induced motor neurons. Notably, the trends we observed in the hyperproliferative population were diminished when examining the non-hyperproliferative cells, highlighting the utility of isolating populations with inherently different reprogramming propensities (Fig. S3D).

Using the fluorescent mRuby2 reporter to identify cells expressing the transgenes, we calculated the reprogramming efficiency for each transcription factor reporter. We define efficiency as the percent of reprogrammed cells normalized to the number of cells expressing the delivered genetic cargo. Reprogramming purity and efficiency are similar for a given transcription factor encoding (Fig. 3E-G). We achieve high efficiency as 37-50% of all transgene-expressing cells convert to iMNs.

### A compact, single-transcript cassette of transcription factors increases conversion

To allow for simple, uniform transcriptional control and ensure co-delivery, we aimed to identify the optimal encoding of all three transcription factors onto a single transcript. Putatively, the ordering of genes influences gene expression. Genes at upstream positions are expected to be translated at greater frequencies than at downstream positions (Mizuguchi et al., 2000). However, relative levels of expression from polycistronic cassettes are context- and encoding-dependent (Carey et al., 2011; Kim et al., 2011; Liu et al., 2017; Rosa et al., 2018; Wang Li et al., 2015; Zhu et al., 2023). Thus, while we had identified putatively optimal profiles of ectopic expression for the three transcription factors, we hypothesized that the ordering on a polycistronic cassette may not fully prescribe expression levels. Therefore, we sought to identify the optimal encoding of all three factors by exploring all permutations on a single transcript, polycistronic cassette. In addition, through screening a variety of C-terminal tags, we identified a tagged Ngn2 variant that does not reduce reprogramming and allows us to identity ectopic Ngn2 via immunostaining (Fig. S4A-F). We compared our original NIL cassette with the untagged Ngn2 and tagged Ngn2^x3HA^. Surprisingly, replacement of Ngn2 with Ngn2^x3HA^ in the NIL cassette increases yield and purity (Fig. 4A, S4G).

**Fig. 4.**
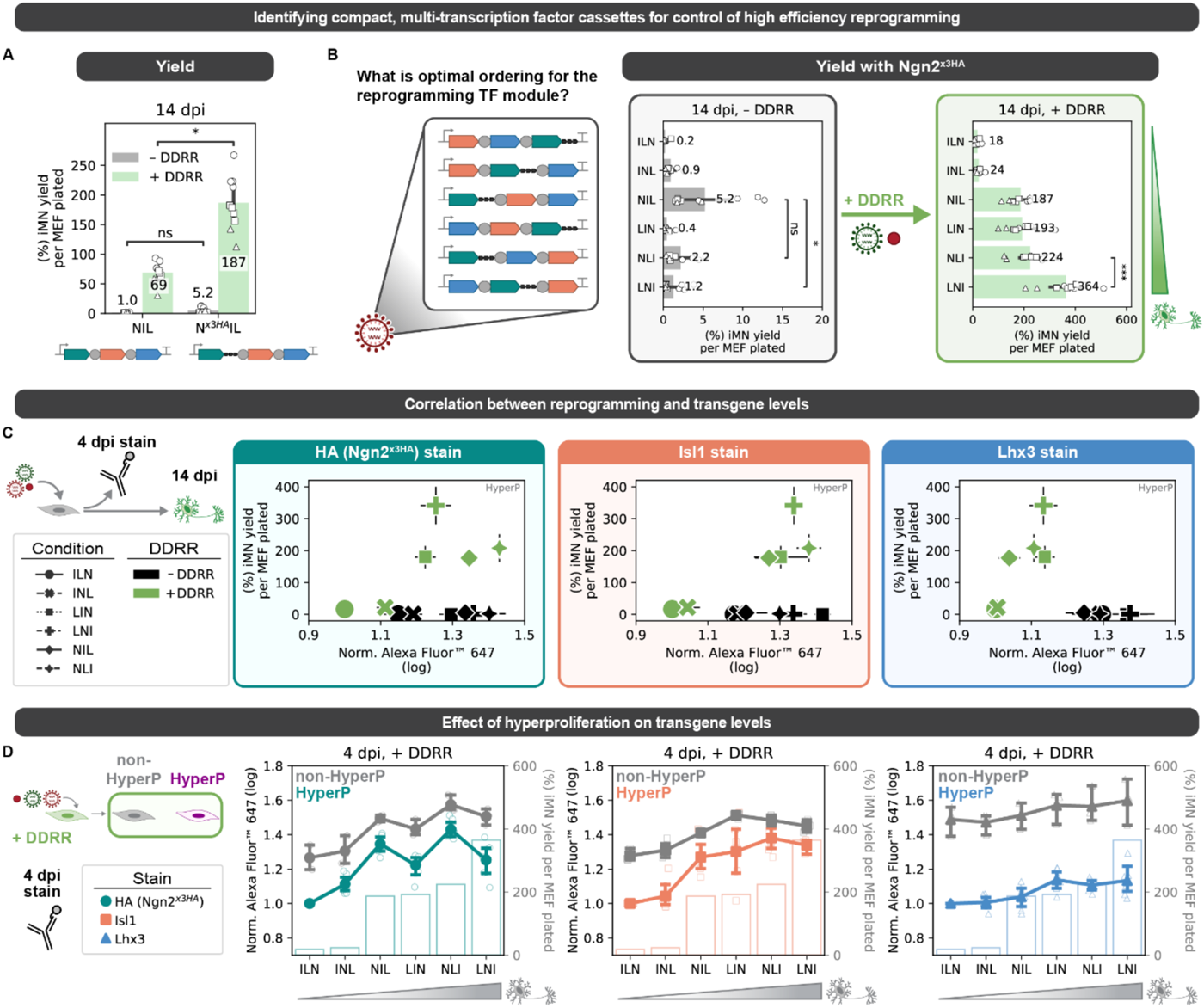
A compact, single-transcript cassette of transcription factors increases conversion. A. Reprogramming yield at 14 dpi for the polycistronic NIL cassette ± DDRR for the untagged Ngn2 and the tagged Ngn2^x3HA^. B. Reprogramming yield at 14 dpi for the different polycistronic transcription factor cassettes without DDRR (left) vs. with +DDRR (right) in order of increasing reprogramming events with DDRR. All cassettes contain the tagged Ngn2^x3HA^. Mean is shown with 95% confidence interval; marker styles denote biological reps; n = 3 biological reps per condition; one-tailed t-test with Bonferroni correction. C. Reprogramming yield at 14 dpi vs. immunofluorescence at 4 dpi for the different polycistronic cassettes without DDRR (black) and with +DDRR (green). Mean is shown with ± standard error of mean (s.e.m.); n = 3-6 biological reps per condition; markers denote reprogramming conditions. Immunofluorescence is normalized such that the mean log expression for ILN + DDRR is 1 in each biological replicate. D. Transcription factor immunofluorescence at 4 dpi for reprogramming with DDRR in non-hyperproliferative (grey) vs. hyperproliferative cells (colored) for HA/Ngn2^x3HA^ (left), Isl1 (middle), and Lhx3 (right). Polycistronic cassettes are ordered by increasing reprogramming events with DDRR (reprogramming yields with DDRR from Fig. 4B are overlaid as bar charts for reference). Mean with biological reps are shown with ± standard error of mean (s.e.m.); n = 3-6 biological reps per condition; marker and color denote antibody used. Significance summary: p > 0.05 (ns); *p ≤ 0.05; **p ≤ 0.01; ***p ≤ 0.001; and ****p ≤ 0.0001

With our set of three transcription factors detectable by immunofluorescence (Ngn2^x3HA^, Isl1, and Lhx3) (Fig S3F), we next built all six permutations of the three factors in polycistronic cassettes using Ngn2^x3HA^, Isl1, and Lhx3 separated by 2A tags (Fig. 4B, S4H). Addition of DDRR resulted in different trends in reprogramming outcomes across the cassettes. In the absence of DDRR, placement of the pioneer transcription factor Ngn2^x3HA^ upstream generates the highest yield and purity (i.e. NIL (Ngn2^x3HA^-2A-Isl1-2A-Lhx3) and NLI (Ngn2^x3HA^-2A-Lhx3-2A-Isl1)). However, with DDRR, LNI (Lhx3-2A-Ngn2^x3HA^-2A-Isl1) reprograms at the highest rate to achieve 50% purity and 360% yield (Fig. 4B, S4H). We did not observe cassette-specific differences in the percent of hyperproliferative cells (Fig. S4I), suggesting that differences in the designs do not alter reprogramming by changing the rate of proliferation.

We hypothesized that differences in the levels of transcription factors may explain differences in the reprogramming rates between the various encodings. To measure transcription factor levels, we probed the levels of each transcription factor in single cells using immunofluorescence and quantified via flow cytometry. For the detection of Ngn2^x3HA^, we could directly measure the transgenic protein via the HA tag. However, for Isl1 and Lhx3 staining, we measure total levels of protein including endogenous and ectopic protein expression. We observed that each cassette design led to a range of expression levels across the bulk population (Fig. S4J).

As expected, the inclusion of the DDRR module increases proliferation and decreases transcription factor levels in the hyperproliferative cells for some cassette designs (Fig. 4C, S4J). To examine proliferation-mediated and cassette-specific effects on transcription factor levels, we looked at immunofluorescence in different reprogramming subpopulations for Ngn2, Isl1, and Lhx3 (Fig. 4D). The most effective designs showed higher levels of Ngn2 expression, whereas less effective cassettes had low Ngn2 expression (Fig. 4C-D). When controlling for the presence of DDRR, expression of Lhx3 was broadly reduced in hyperproliferative cells while Ngn2 and Isl1 were minimally affected (Fig. 4D).

Based on the individual factor titration, we expected cassettes with the highest Ngn2^x3HA^ and lowest Lhx3 levels to reprogram the at the highest rate. However, the best performing design, LNI, does not have the highest Ngn2 nor the lowest Lhx3 levels (Fig. 4C-D). Together, these results demonstrate that the encoding of transcription factors in polycistronic cassettes influences the expression of factors and this influence is independent of differences in proliferation history (S4I). For small design spaces, functional screening of single-transcript cassettes of transcription factors remains the best approach for optimization.

### Driving hyperproliferation improves direct conversion of adult human fibroblasts

Having identified a compact cocktail that reduces variability and increases the yield of motor neurons from murine cells, we wondered how these cocktails might impact reprogramming of primary adult human dermal fibroblasts, where yields and maturity remain limited (Fig. 5A). We hypothesized that differences in proliferation rates between MEFs and human fibroblasts may account for the difference in reprogramming yield. Thus, we focused on increasing the proliferation rate of adult human fibroblasts by addition of genes known to drive proliferation. We quantified proliferation using a CellTrace dilution assay measured at 7 dpi (Fig. 5B).

**Fig. 5.**
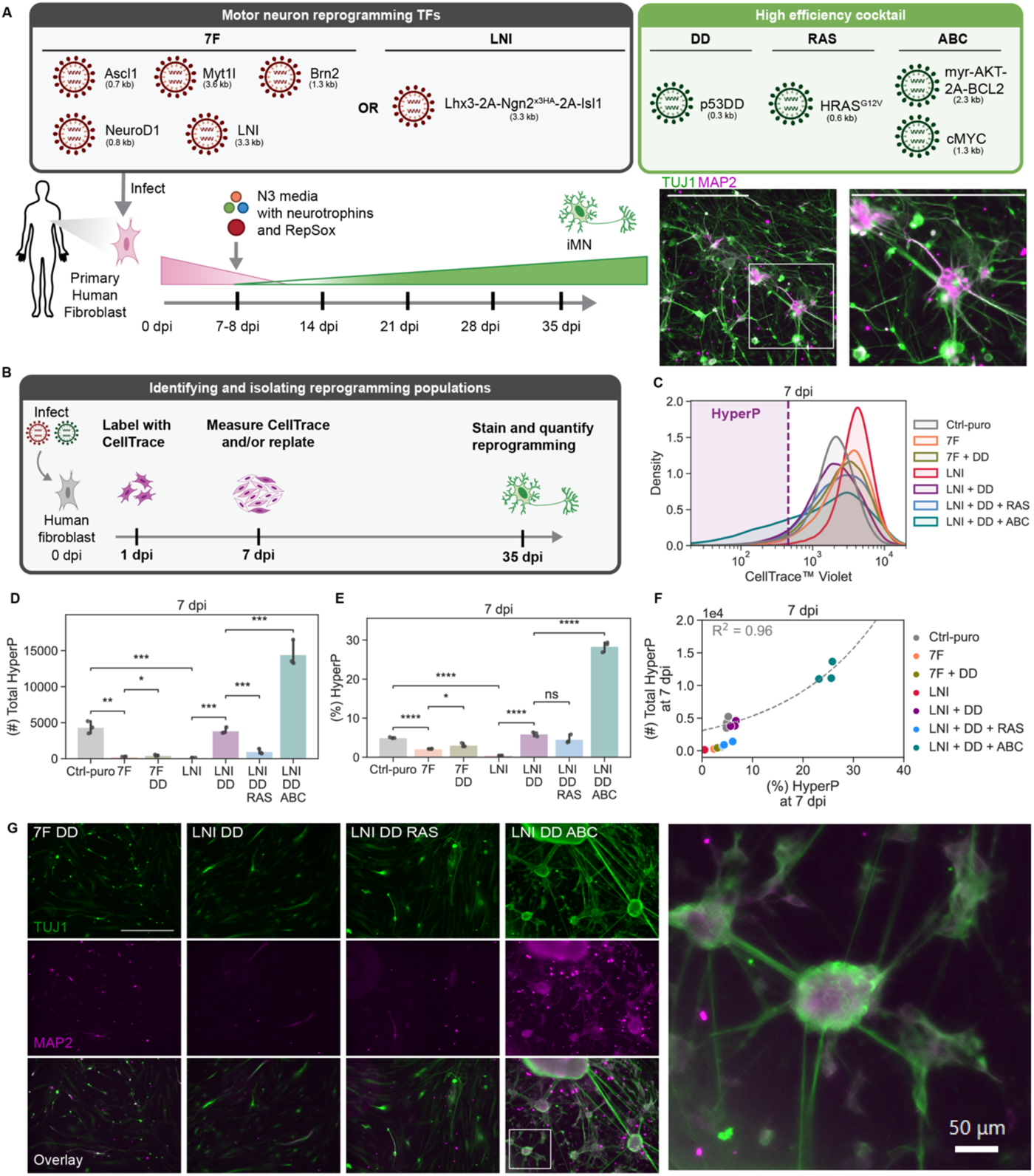
Driving hyperproliferation improves direct conversion of adult human fibroblasts. A. Schematic depicting the reprogramming process for human dermal fibroblasts. B. Schematic of the CellTrace dilution assay. HDFs are labeled 1 dpi and their fluorescent intensity is measured at 7 dpi, where cells that have gone through many divisions have low fluorescence. C. Representative CellTrace distributions across conditions. Hyperproliferative cells are defined relative to the 5%-lowest fluorescent cells in a control puro (Ctrl-puro) infection condition for a given biological replicate. D. Hyperproliferative (HyperP) percent at 7 dpi across conditions. Ctrl-puro is 5% HyperP by definition. Mean is shown with 95% confidence interval; n=1 biological reps per condition; One-tailed t-test with Bonferroni correction. E. Hyperproliferative (HyperP) total number at 7 dpi across conditions. Mean is shown with 95% confidence interval; n=1 biological reps per condition; One-tailed t-test with Bonferroni correction. F. HyperP total number vs. percent at 7 dpi across all reprogramming conditions. An exponential fit is shown. G. Representative images at 35 dpi of cells fixed and stained with TUJ1 and MAP2. Scalebar represents 500 µm.

As expected, the inclusion of p53DD increases hyperproliferation and the total number of cells that undergo a period of hyperproliferation in both the 7F (BAM + NIL + NeuroD1) and LNI conditions (Fig. 5C-F). As expected from comparing 6F and NIL in murine cells, LNI + DD achieve a greater number and fraction of hyperproliferative cells compared to 7F + DD (Fig. 5D-E). Unlike in the murine system, the addition of HRAS^G12V^ reduces the total number of hyperproliferative cells (Fig. 5D) to lower than expected based on the fraction of hyperproliferative cells (Fig. 5F). To increase the total number of cells that have undergone a period of hyperproliferation, we introduced myr-AKT, BCL2, and c-MYC (ABC), which are known to drive proliferation in oncogenesis (Park et al., 2018). Addition of ABC to adult human fibroblasts resulted in a three-fold increase in the number of hyperproliferative cells co-transduced with LNI + DD (Fig. 5D).

With the human-specific module to drive proliferation, we assessed neuron formation induced by the different cocktails by immunofluorescence staining for the neuronal markers TUJ1 and MAP2. In alignment with the increase in proliferation rates, addition of ABC to NIL, LNI, and 7F generates more TUJ1 and MAP2-positive cells (Fig 5G, S5A). Cells reprogrammed with the addition of ABC display mature, neuronal morphologies and cultures form clusters of neurons with interconnecting neurites (Fig 5G, S5A). As expected from the reduction in hyperproliferative cells, inclusion of HRAS^G12V^ reduces the formation of human iMNs. Conversely, the introduction of DD and ABC increases proliferation and reprogramming rates. Additionally, in the presence of DD+ABC, we observed that higher seeding densities increase the fraction of hyperproliferative cells (Fig. S5B-D). By MAP2 and TUJ1 staining at 35 dpi, we observed similarly dense and connected networks of morphologically mature neurons with LNI+DD+ABC (Fig. S5E). As direct conversion of primary human fibroblasts offers a powerful platform for disease modeling and drug discovery, improved rates of reprogramming are essential for utilizing these reprogrammed surrogate cells at scale for screening.

### Compact cocktail enables neurotrophic factor-free reprogramming

During *in vitro* direct conversion, transcription factors direct cell fate in combination with external signaling cues from growth factors and other biochemicals. Conversion protocols often rely on specific cocktails of small-molecules and growth factors (Babos et al., 2019; Son et al., 2011). We wondered how media supplements might impact the direct conversion process. To explore how media supplements affect our reprogramming efficiency, we reprogrammed cells with and without neurotrophic factors and the small molecule TGFβ-inhibitor, RepSox, which is added in our high-efficiency DDRR module. In the absence of neurotrophic factors, NIL-only induced direct conversion does not generate motor neurons (Fig. 6A, S6A). However, with the high-efficiency DDRR module, the reprogramming yield of Hb9::GFP positive cells remains high in the absence of neurotrophins and RepSox. However, in the absence of both, Hb9::GFP+ cells adopt more fibroblast-like morphologies (Fig. 6B). Together these data suggest that neurotrophic factors and RepSox are dispensable for conversion but may support neuronal maturation via morphological remodeling.

**Fig. 6.**
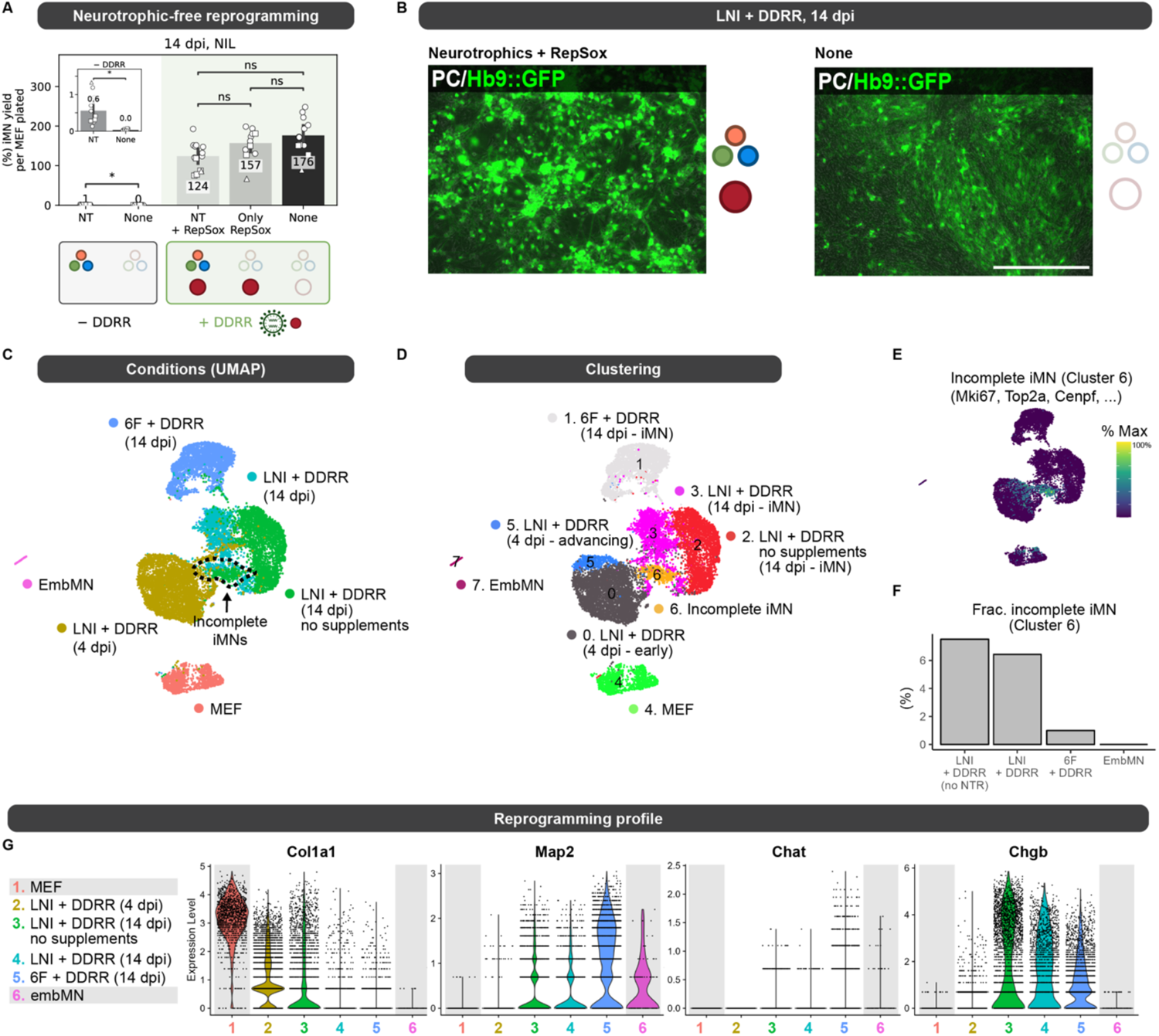
Compact cocktail enables neurotrophic-free reprogramming. A. Reprogramming yield at 14 dpi for NIL ± DDRR. NIL infected cells were reprogrammed in N3 media ± neurotrophic growth factors (NT). NIL + DDRR infected cells were reprogrammed in N3 media with both NT and RepSox, with only RepSox, and with neither. Inset shows iMN yield for NIL (without DDRR) with zoomed-in axis for clarity. Mean is shown with 95% confidence interval; marker styles denotes biological reps; n = 3 biological reps per condition; two-tailed t-test with Bonferroni correction. B. Images of cells at 14 dpi reprogrammed by LNI with the Ngn2^x3HA^ tag + DDRR, in N3 media containing neurotrophic growth factors and RepSox (left) and with no supplements (right). Scale bar represents 500 µm. C. UMAP of different reprogramming conditions. Embryonic motor neurons (EmbMN, dark grey) and mouse embryonic fibrobalsts (MEFs, light grey) serve as end and initial cell identities references. LNI is the polycistronic cassette with Ngn2^x3HA^. EmbMN were isolated from a primary Hb9::GFP+ spinal cord at E12.5 and bioinformatically isolated based on Isl1, Lhx3, and Mnx1 expression (n=91). All 14 dpi conditions were sorted for Hb9::GFP+ cells prior to loading for scRNA-seq, and LNI + DDRR 4 dpi was sorted for hyperP cells only. D. Clustering of UMAP from Fig. 5C. Annotated clusters were determined by looking at overlap with reprogramming conditions (Fig. 5C-D, S5B) and by looking at cluster biomarkers. E. Incomplete reprogrammed motor neurons revealed by clustering. Mki67 expression is shown but other genes expressed during G2/M (e.g. Top2a, Cenpf) show similar trends. F. Percentage of cells in the incomplete iMN cluster 6 across reprogramming conditions at 14 dpi with EmbMN as a baseline. G. Violin plots of scRNA expression of stereotypical markers of MEFs (Col1a1) and motor neuron maturation (Map2, Chat, Chgb) for reprogramming conditions after incomplete iMN cluster 6 is removed for representative genes of motor neuron identity. Significance summary: p > 0.05 (ns); *p ≤ 0.05; **p ≤ 0.01; ***p ≤ 0.001; and ****p ≤ 0.0001

To examine the transcriptional states of our reprogrammed cells, we generated and sequenced single-cell RNA sequencing libraries of cells collected across the conversion timeline and in the absence of media supplements. We induced reprogramming using the optimized polycistronic LNI cassette with the tagged Ngn2^x3HA^ and the high-efficiency DDRR module, and collected Hb9::GFP+ cells via fluorescence-activated cell sorting (FACS) at 14 dpi. With this cocktail, we collected cells reprogrammed either with or without media supplements. We also collected hyperproliferative cells at 4 dpi to capture early transcriptional states in the reprogramming process. We used MEFs and embryonic motor neurons (embMN) as reference cell states for the beginning and target cell types (Fig. 6C). In the hyperproliferative cells, we identified a cluster with increased expression of transcription factors involved in neurogenesis, including Bhlhe22 and Neurod1 (Tutukova et al., 2021), suggesting these cells may be advancing toward iMNs (Fig. 6D, S6C). Transcription factors express at higher levels in the “advancing” state. These data are consistent with a model in which high expression of the transcription factors drive hyperproliferative cells to convert at high rates.

Considering Hb9::GFP positive cells at 14 dpi, we were able to identify a cluster of incompletely reprogrammed cells (Fig. 6C,D). Although Hb9::GFP positive, these cells are enriched for markers of proliferation (e.g. Mki67) and appear as a small fraction of cells across cocktails (Fig. 6E-F, S6B). The higher percent of incomplete iMNs in LNI + DDRR compared to 6F + DDRR is consistent with our observation that the three transcription factor cocktail expands the population and window of time over which cells proliferate and reprogram (Fig. 2F, S2K, 6F).

To compare different cocktails, we focused on the fully reprogrammed populations of motor neurons. Across the different reprogramming cocktails, we observe differences in markers of maturity (Fig 6G). Differences in markers of neuronal maturity between iMNs and embMNs may be attributed to the early developmental stage from which the primary embMNs were isolated (13.5 dpc). As expected from the observed morphologies (Fig 6B), excluding media supplements in LNI + DDRR generates cells with elevated expression of the fibroblast-specific matrix protein Col1a1. Expression of the transcription factors, Ngn2 and Lhx3, does not significantly differ between the reprogrammed iMNs and embMN, whereas Isl1 expression is lower in iMNs compared to embMNs (Fig. S6D). Together, these observations demonstrate that the motor neuron transcription factors are not expressed at supraphysiological levels in reprogrammed cells. Moreover, induced motor neurons cluster primarily by cocktail, indicating that cells vary as a function of the reprogramming cocktail (Fig. S6B).

### Compact cocktail generates engraftable neurons

In generating large numbers of induced motor neurons at a scale relevant for cell-based therapies (Ceto et al., 2020), we wondered if reprogrammed cells could survive when implanted into the central nervous system. Replacement of damaged or diseased tissues and organs represents a major goal of cellular reprogramming. Potentially, autologous cell grafts generated from accessible cell donor populations could be used to restore lost populations of cells and support renewed function. However, low rates of reprogramming limit the potential for central nervous system (CNS) applications. Additionally, while cells may offer therapeutic effects, implanted cells may not survive engraftment (Tabar and Studer, 2014). Since the high-efficiency DDRR module supports the generation of motor neurons by direct conversion at scales sufficient for grafting (e.g. millions of cells, Fig S7A) even in absence of media supplements, we next evaluated *in vivo* transplantation outcomes for iMNs reprogrammed via LNI + DDRR. To examine grafting, we transplanted iMNs in healthy murine brains as well as animal subjected to a small acute N5-(1-Iminoethyl)-L-ornithine (L-NIO) induced stroke-injury (Fig. 7A). The striatum was chosen for initial proof-of-principle for iMN grafting because it is a readily accessible anatomic site that allows for reproducible injuries and transplantation injections (O’Shea et al., 2022). Additionally, the striatum contains neural tissue with both neuronal cells bodies and myelinated axon bundles as well as a diversity of glia to enable assessment of graft-host tissue integration (O’Shea et al., 2022).

**Fig. 7.**
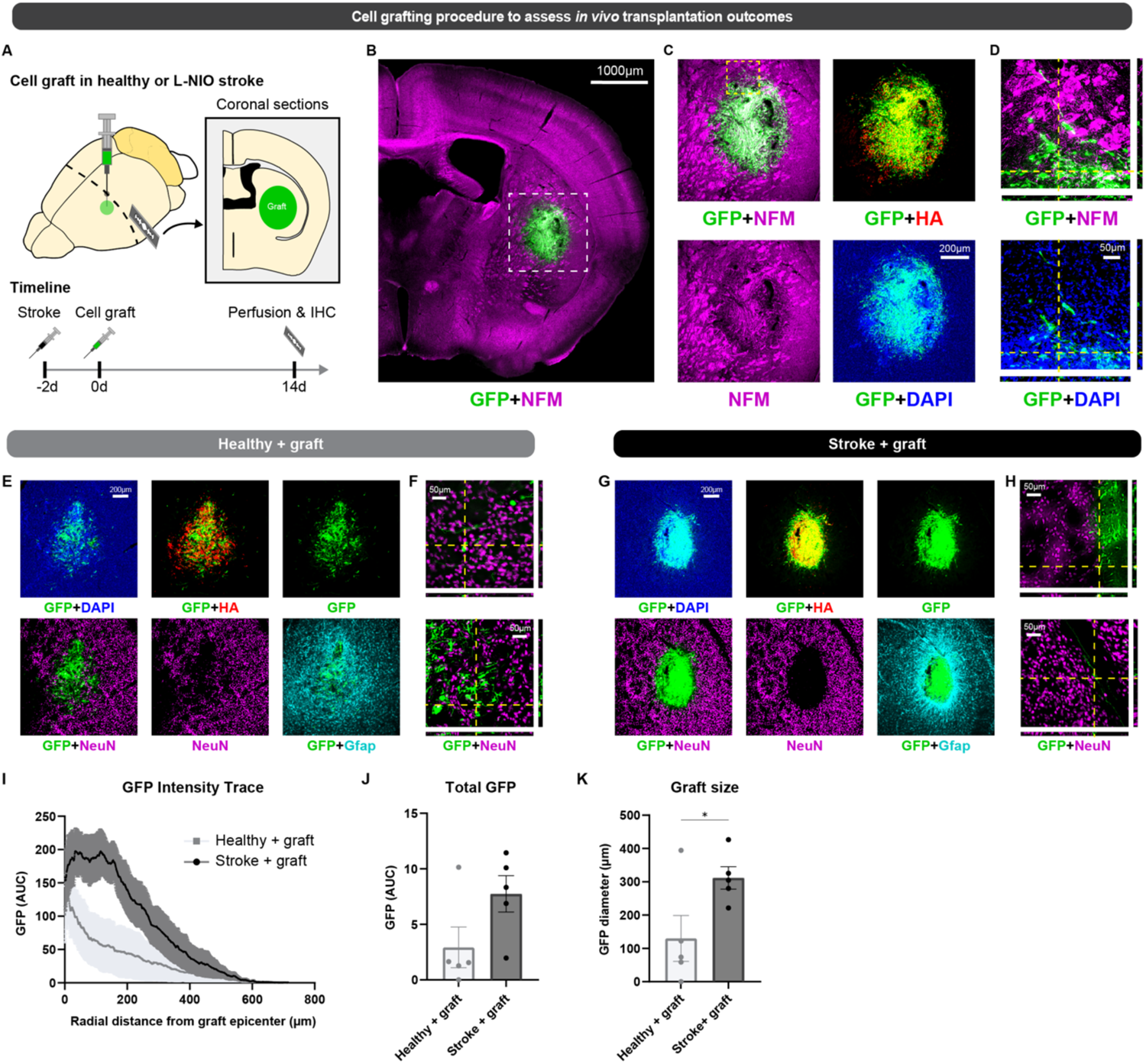
Compact cocktail generates engraftable neurons. A. Schematic outlining iMN grafting into healthy or stroke murine brains. Striatal strokes were induced two days prior to cell grafting by local injection of L-NIO. iMN graft survival correlates with time of grafting (Fig. S7B). B-D. Survey and detailed immunohistochemistry images showing integration of GFP/HA-positive iMNs with striatal stroke lesions. iMNs express neuronal marker, neurofilament (NFM). E-H. Coronal sections healthy (E-F) and stroke-induced (G-H) murine striatums two weeks after reprogrammed iMN cell graft procedure at various objectives. Sections were imaged for GFP, neurofilament (NFM), NeuN, and Gfap (glial fibrillary acidic protein). G. Trace of GFP intensity from graft epicenter radially in graft+stroke and graft only mouse striatum. H. Quantification of total intensity of GFP-positive grafted iMNs at graft injection site. Quantified by integrating GFP values up at 350 microns I. Graft size defined at 60th-90th percentile of GFP intensity radially. Significance summary: p > 0.05 (ns); and *p ≤ 0.05

At two weeks post grafting, we detected numerous GFP-positive and HA-positive iMNs at the striatal injection site across both healthy and stroke conditions (Fig. 7B-H, S7C-D). iMN grafts robustly expressed neurofilament proteins but not the mature neuronal marker, NeuN (Fig. 7E-H). iMNs were negative for the striatal medium spiny neuron marker Darpp-32 suggesting that reprogrammed cells maintained their motor neuron identity *in vivo*, even in neural tissue niches where they do not naturally reside (Fig. S7E-F). While most iMNs remained local to the injection site, grafts integrated with host tissue and extended axonal projections several hundred microns into healthy neural tissue (Fig. 7F, S7D). There was improved graft survival at small L-NIO strokes compared to healthy tissue, which was consistent with results from previous grafting studies using neural progenitor cells (Fig. 7I-K) (Adewumi et al., 2023). These data show that our reprogramming protocol can directly convert donor cells at sufficient numbers necessary for CNS grafting and that grafted iMNs survive, maintain their neuronal identity, and integrate with neural tissue in the murine brain.

## Discussion

Despite improved techniques and tools, low reprogramming rates and diverse sources of extrinsic variation limit our ability to interrogate and engineer cell-fate transitions. To reduce extrinsic variation and resolve the impact of transcription factor levels on reprogramming, we reduced the number of viruses to two by developing a minimal motor neuron transcription factor module and a compact high-efficiency module based on our previously developed chemo-genetic DDRR cocktail. With the statistical power conferred by higher reprogramming rates, we were able to identify how expression of individual transcription factors impacts reprogramming. Through systematic titration, we demonstrate that MEF-to-iMN direct conversion efficiency positively correlates with levels of the pioneer transcription factor Ngn2 early in the reprogramming process. To improve the performance and controllability of our reprogramming cocktail, we used screening to identify the optimal encoding of transcription factors within a single transcript. Using only two viruses, we increase the reprogramming rate 100-fold over the original high-efficiency cocktail encoded on eight viruses. Our improved high-efficiency cocktail generates motor neurons at scale (e.g. millions) via direct conversion. Additionally, we applied our observations to develop compact reprogramming cassettes that improved direct conversion rates in human cells and allowed *in vivo* grafting of induced motor neurons into the murine central nervous system.

While transcription factors drive cell-fate reprogramming, processes such as transcription and proliferation impact expression of transgenic transcription factors and reprogramming. We set out to resolve how cells integrate contradictory cues from proliferation and transcription factor expression during successful reprogramming events. We find that hyperproliferative cells reprogram at higher rates despite reduced levels of transgene expression. Our results suggest that proliferation confers a state of plasticity that exceeds what high expression of transcription factors alone can confer. Once cells attain this proliferation-mediated state of plasticity, transcription factor levels direct cells in distinct ways. High levels of the pioneer transcription factor Ngn2 correlate to higher rates of reprogramming, whereas expression of Isl1 and Lhx3 do not show positive correlations early in reprogramming. Putatively, early opening of chromatin via pioneer activity of Ngn2 may provide accessibility to the other transcription factors. Additionally, high rates of transcription in hyperproliferative cells allow cells to reprogram at near-deterministic rates (Babos et al., 2019). Potentially, the levels of Ngn2 may drive high rates of transcription in hyperproliferative cells and enable cells to reprogram at high rates.

By isolating the hyperproliferative cells in our analyses, we could deconvolute proliferation as a confounding factor to examine how transcription factors influence reprogramming from this privileged cell state. Specifically, we found that Brn2 alone reduces proliferation and reprogramming, whereas Ascl1 and Myt1l can increase reprogramming without changing proliferation rates. Further, eliminating Ascl1 from the reprogramming cocktail may remove differences in reprogramming due to distinct mechanistic pathways of the pro-neural, pioneering transcription factors, Ascl1 and Ngn2 (Aydin et al., 2019; Vainorius et al., 2023). Potentially, variation in Brn2 expression in 6F cocktails may explain previously observed variability in reprogramming rates as well as observations by others that Brn2 cocktails rapidly drive cells to exit cell cycle (Vierbuchen et al., 2010; Wapinski et al., 2013). Notably, the transcription factor-specific trends we observed in the hyperproliferative population were diminished or eliminated in the non-hyperproliferative cells (Fig S3D). As hyperproliferative cells often represent a small fraction of reprogramming cultures, trends observed in bulk populations may mask the processes that actually direct reprogramming within these highly reprogrammable cells.

Reprogramming has been extensively studied through transcriptional profiling (Cahan et al., 2014; Cates et al., 2020; Jindal et al., 2023; Mall et al., 2017; Nair et al., 2023; Schiebinger et al., 2019; Zhao et al., 2018; Zviran et al., 2019). However, transcription factors activate gene regulatory networks as proteins. Post-translational regulation of protein levels can generate large variance between levels of RNA and protein for the same gene (Hafner et al., 2017; Ideker et al., 2001; Lundberg et al., 2010). Here we focused on measuring protein levels to examine how transgenic transcription factors influence reprogramming events. In our individual transcription factor titration, we quantified the expression of the ectopic transgenes. For our polycistronic cassettes, we captured the combined expression of ectopic and endogenous of Isl1 and Lhx3 via immunostaining. As we tagged Ngn2 (Ngn2^x3HA^), we could specifically identify the transgene via the HA tag. For both types of measurements, we could map how levels of transcription factors translate to rates of reprogramming, suggesting the potential for designing optimal levels and stoichiometries of factors. While we expect that the total levels of transcription factors are what dictate reprogramming, the ectopic factors represent the primary species for driving and guiding cell-fate transitions.

Although we have gained insight into how transcription factor levels and proliferation drive reprogramming, there are still open questions. To optimally guide cells through transitions, we sought to understand how ectopic levels of individual transcription factors impact reprogramming. However, in designing single-transcript cassettes, we could not predict which designs would perform best. While high expression of Ngn2 at 4 dpi differentiated effective from non-effective designs, the ordering was not fully predictive of expression. Thus, we conclude that screening provides the best tool to identify optimal designs as the strict rules of design of polycistronic cassettes remain unclear. Potentially, differences in expression of factors that emerge later in the conversion timeline will impact a single cell’s trajectory. Longitudinal tracking of Oct4 in reprogramming human fibroblasts indicates that cells that establish high expression early sustain expression over the course of reprogramming (Ilia et al., 2023). Future systems that can more precisely vary and measure all three ectopic transcription factor levels in single cells may help elucidate how cells interpret transcription factor combinations to acquire new cell fates.

In profiling the transcriptomes of induced motor neurons generated from different cocktails, we found that our optimized, two-virus cocktail leads to a higher fraction of incompletely reprogrammed cells compared to our six-factor cocktail. Eliminating these partially converted cells may be important in developing *in vitro* disease models and cell-based therapies (Locatelli et al., 2022). Alternatively, given the power of the transcription factors to drive cells to a post-mitotic fate in the presence of the oncogenes, cocktails of neuronal transcription factors may be useful in arresting growth of transformed cells with minimal impact on non-proliferative cells.

While cell lines engineered with inducible transcription factors provide a system for limiting the extrinsic variance associated with viral infection, these systems provide limited flexibility in modifying transcription factor cocktails (Elbaz et al., 2022; Jain et al., 2023). Additionally, achieving rapid, scalable reprogramming *in vitro* and translatable technologies for *in vivo* reprogramming requires efficient transgenic systems that induce reprogramming without the need for extensive cell line engineering. We aimed to develop systems that could be translated to *in vivo* reprogramming and to adult human fibroblasts from diverse donors. By developing a species-specific high-efficiency module, we increase proliferation in human fibroblasts which correlates to higher reprogramming rates and more morphologically mature motor neurons. Unlike iPSC-derived neurons, directly converted neurons retain signatures of aging (Mertens et al., 2015). Developing scalable methods for generating neurons from primary adult fibroblasts will support disease modeling in adult-onset neurodegenerative disease such as Amyotrophic lateral sclerosis, Parkinson’s disease, Huntington’s disease, Alzheimer’s disease, and Frontotemporal dementia. By generating neurons from a greater number of patient-derived cells, direct conversion may provide a more representative survey of cellular diversity within a single patient and avoid clonal bias introduced in clone-based iPSC-derived models.

With our optimized cocktail, we achieve neurotrophic factor-free motor neuron reprogramming and show that our reprogrammed motor neurons survive *in vivo* grafting. As the *in vivo* environment may provide varying or limited neurotrophic support, our work highlights that cells can reprogram in the absence of neurotrophic support. Moreover, reprogrammed cells can be grafted following focal injury as needed for the replacement of damaged tissue, opening future studies for *in vivo* reprogramming and cellular therapies.

Collectively, our results show that proliferation history and transcription factor expression combine to drive cell-fate transitions. By developing tools that more precisely perturb the reprogramming process, we will improve our understanding and control of cellular reprogramming. Further, by developing single transcript designs, we open the potential for integrating control of reprogramming with the expanding array of synthetic biology tools and gene circuits for tailoring expression within specific subpopulations. Beyond targeting specific cell types, control of transgenes to induce reprograming of human cells will improve safety and efficacy of gene and cell-based therapies for regeneration of neural tissue.

## EXPERIMENTAL MODEL AND SUBJECT DETAILS

### Cell lines and tissue culture

HEK293T, Plat-E, and mouse embryonic fibroblasts were cultured in DMEM supplemented with 10% FBS at 37°C, 5% CO2. Plat-E cells were selected using 10 µg/mL blastocidin and 1 µg/mL puromycin every three passages. Human adult dermal fibroblasts were cultured in DMEM supplemented with 15% FBS and non-essential amino acids at 37°C, 5% CO2. The following are sex of primary human fibroblasts used in this study: female (GM05116). Cultures were periodically tested for mycoplasma.

### MEF dissection and isolation

C57BL/6 mice were mated with mice bearing the Hb9::GFP reporter. Mouse embryonic fibroblasts were isolated at E12.5-E14.5 under a dissection microscope. Embryos were sorted into non-transgenic and Hb9::GFP+ by using a blue laser to illuminate the spinal cord to identify the presence of Hb9::GFP+ cells. After removing the head and internal organs to avoid contaminating neurons and other cells, razors were used to break up the tissue for 5 minutes in the presence of 0.25% Trypsin-EDTA. Up to two embryos were processed at the same time. The preliminary suspension was neutralized, resuspended, and triturated with 0.25% Trypsin-EDTA. Again, the suspension was neutralized, resuspended, and filtered through a 40 µm cell strainer. MEF cultures were expanded on 0.1% gelatin coated 10 cm dishes, using one 10 cm dish per embryo. MEFs were expanded until ∼80% confluent, passaged, and expanded for at least 3-4 days. Passage 1 MEFs were tested for mycoplasma, cryopreserved in 90% FBS and 10% DMSO, and kept in liquid nitrogen.

### Animals

All *in vivo* experiments involving the use of mice were conducted according to a protocol approved by the Boston University IACUC (Protocol Number: PROTO202000045). Female C57BL/6 mice (RRID:IMSR_JAX:000664) were used for all *in vivo* experiments and underwent surgical procedures at 8 weeks of age. All experiments involving mice were conducted within approved Boston university facilities and mice were housed in a 12-hour light/dark cycle in a for-purpose facility with controlled temperature and humidity. Mice were provided with food and water ad libitum. Mice were administered post-surgical analgesia (buprenorphine, 0.1mg/kg) for 2 days after each surgery. No animals in the study that received cell grafts were administered with any immunosuppression drugs.

## METHOD DETAILS

### Plasmid construction

Entry vectors were constructed by Gibson and Golden gate cloning. Retroviral vectors were constructed by Gateway cloning into pMXs-WPRE-DEST using an LR reaction, respectively. Viral plasmids were confirmed via Sanger or whole plasmid sequencing. For a complete list of plasmids, see supplementary files.

### Viral transduction and iMN reprogramming of MEFs

Plat-E cells were seeded at 850k per 6-well onto 6-well plates coated with 0.1% gelatin for at least 10 minutes at room temperature one day prior. The next day, ∼80% confluent Plat-E cells were transfected with 1.8 µg of DNA per well using a 4-5:1 ratio of µg PEI:µg DNA. The next day, a media change was performed with 1.25 mL fresh DMEM supplemented with 10% FBS and 25 mM HEPES to help buffer. MEFs were also seeded at 10k per 96-well onto 96-well plates coated with 0.1% gelatin. The next day, viral supernatant was collected, filtered through a 0.45 µm filter, and 1.25 mL fresh DMEM supplemented with 10% FBS and 25 mM HEPES was added again to the Plat-E cells. MEFs were transduced with 11 µL of viral supernatant per viral cassette. Fresh DMEM supplemented with 10% FBS was added to reach a final volume of 100 µL per 96-well. 5 µg/mL polybrene was supplemented to increase transduction efficiency. Transduction was repeated for a second day. One day after the second viral transduction was considered 1 dpi and 100 µL per 96-well fresh media was swapped in. At 3 dpi, media was switched to N3 media (DMEM/F-12 supplemented with N2, B27, and neurotrophic growth factors, BDNF, CNTF, GDNF, FGF all at 10 ng/mL) (Babos et al., 2019; Chanda et al., 2014). For conditions with RepSox, 7.5 µM RepSox was also supplemented into the N3 media starting at 3 dpi. For experiments needing larger well sizes, MEF seeding and viral supernatant volumes were scaled accordingly by well surface area.

### Quantification of reprogramming yield, purity, and efficiency

Reprogrammed cells were dissociated using 0.025% Trypsin-EDTA prior to 6 dpi and DNase/papain from 6 dpi onwards. One vial of papain (>100 U/vial) and one vial DNase (>1,000 U/vial) were dissolved in 6 mL of DMEM/F-12. 50 µL of this dissociation media was used per 96-well. Cells were incubated with dissociation media at 37°C, 5% CO2 for ∼15 minutes or until starting to detach. 100 µL of media was added to neutralize the reaction, then cells were resuspended in 300 µL PBS. All cells in a well were recorded using flow cytometry to detect Hb9::eGFP fluorescence. iMNs were defined by gating the brightest Hb9::GFP fluorescent cells (Fig. S1A) (Babos et al., 2019). Yield was calculated by dividing the total number of Hb9::GFP positive cells detected via flow by the total number of initially seeded cells which was generally 10k/96-well. Purity was calculated by dividing the total number of Hb9::GFP positive cells by the total number of cells detected via flow. Efficiency was calculated by dividing the total number of Hb9::GFP and mRuby2 positive cells by the total number of mRuby2 positive cells detected via flow.

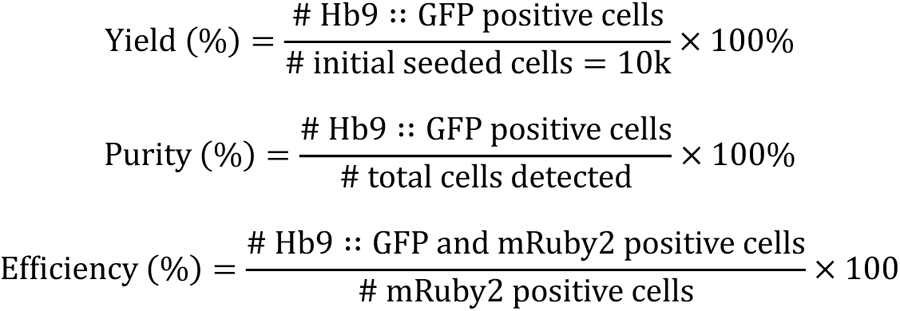

At 1 dpi, cells were first washed with PBS. Then 100 µL/96-well of 1µM CellTrace-Far Red™ or 5µM CellTrace-Violet™ in PBS was added. Cells were incubated at 37°C, 5% CO2 for 30 minutes. Then the staining solution was removed, cells were washed with media to remove residual stain, then fresh media was added. Cells were then cultured or reprogrammed as normal. CellTrace fluorescence was detected by flow cytometry. Cells that undergo a period of hyperproliferation were identified by dilution of CellTrace dye over 72 hours from 1 to 4 dpi for MEF reprogramming and over 144 hours from 1 to 7 dpi for human reprogramming. We denote cells with a history of rapid proliferation at the time point assayed as hyperproliferative (hyperP) and all other cells at 4 dpi as non-hyperP relative to the CellTrace dilution in a control puro infection condition to account for effects on proliferation induced by viral transduction. For MEF and human reprogramming, hyperP cells are defined relative to the 20%-lowest and 5%-lowest fluorescent cells, respectively, in a control puro infection condition for a given biological replicate.

### Flow cytometry and FACS

An Attune™ NxT Acoustic Focusing Cytometer was used for flow cytometry analyzer experiments. A 405 nm laser with a 440/50 filter was used for blue fluorescence (DAPI, CellTrace-Violet™), a 488 nm laser with 510/10 filter was used for green fluorescence (eGFP), a 561 nm laser was used with 585/16 filter for red fluorescence (Alexa Fluor™ 594, mRuby2), and a 638 nm laser with a 670/14 filter was used for far red fluorescence (Alexa Fluor™ 647, CellTrace-Far Red™, Zombie NIR™). FSC-A and SSC-A were used to gate cells, while FSC-H and FSC-A were used to gate singlets using FlowJo. Singlets were exported as csv files to analyze using Python.

FACS was performed using a BD FACSAria. A 488 nm laser with 530/30 filter was used for green fluorescence (eGFP) and a 640 nm laser with a 670/30 filter was used for far red fluorescence (CellTrace-Far Red™, Zombie NIR™). Cells were sorted into N3 media supplemented with penicillin-streptomycin.

For replating experiments, cells were resuspended in their appropriate medias, counted, and replated at 10k/96-well. Penicillin and streptomycin were included for the rest of reprogramming after sorting.

For all cytometry experiments, appropriate controls were included to assess negative populations (*e.g.* un-stained cells, single color controls, secondary antibody only controls, non-targeting antibody controls, etc.).

### Twin reprogramming assays for 4 dpi ectopic transgene reporter and 14 dpi reprogramming

MEFs from a single biological source were seeded in duplicate and reprogrammed using six uORF conditions (five uORFs and one without any uORFs) with the same viral supernatant mixes, such that each twin received the same exact transduction treatments. At 1 dpi, one biological twin was stained with CellTrace-Violet™ to assay at 4 dpi for proliferation history and ectopic transgene expression via the mRuby2 reporter. The other biological twin was kept until 14 dpi to measure reprogramming yield and purity. Each twin was performed with three technical replicates and the results were averaged and plotted against each other for each biological replicate. Thus, each point represents the average geometric mean of mRuby2 expression at 4 dpi across three wells and the average reprogramming rate at 14 dpi across another three wells, for a total of six wells assayed for a single biological replicate of one condition.

### Fixation and immunofluorescent staining

For plate imaging, cells were fixed with 3.7% paraformaldehyde for 1 hour at 4°C. Cells were washed three time with cold PBS and permeabilized with 0.5% Tween/PBS overnight at 4°C. Cells were then blocked with 5% FBS and 0.1% Tween/PBS for 1 hour at 4°C. Cells were then incubated with primary antibody diluted in blocking solution (5% FBS and 0.1% Tween/PBS) overnight at 4°C. Cells were then washed three times with cold 0.1% Tween/PBS, with the third wash being left for at least 20 minutes. Cells were then incubated with secondary antibody diluted in blocking solution (5% FBS and 0.1% Tween/PBS) for 1 hour at 4°C. Cells were then washed three times with cold 0.1% Tween/PBS and 0.1 µg/mL DAPI staining was added for 10 minutes. Cells were washed three more times with cold 0.1% Tween/PBS and imaged. In between incubation steps, the plate was kept covered in foil.

For flow cytometry, cells were dissociated and fixed with 3.7% paraformaldehyde for 15 minutes at room temperature. Cells were permeabilized by 200 µL of 0.5% Tween/PBS for 15 minutes at room temperature, and washed with PBS. 200 µL of primary antibody diluted in blocking solution (5% FBS and 0.1% Tween/PBS) was added and incubated at 4°C for 2 hours with rotation for mixing. Cells were washed three times with 1 mL cold 0.1% Tween/PBS, with the third wash being left for at least 30 minutes at 4°C. Cells were then incubated with 800 µL of secondary antibody diluted in blocking solution for 30 minutes at 4°C and washed three times with 1 mL cold 0.1% Tween/PBS. In between incubation steps, cells were kept covered in foil.

### Viral transduction and iMN reprogramming of human adult dermal fibroblasts

HEK293Ts were seeded at 6.5 million per 10 cm dish onto 10 cm dishes coated with 0.1% gelatin. The next day, each dish of 293Ts was transfected with 10.8 ug pLK-packaging plasmid, 1.2 ug of pHDMG envelope plasmid, and 12 ug of transfer plasmid using a 4-5:1 ratio of ug PEI:μg DNA. 6-8 hours later, a media change was performed with 6.5 mL of 25 mM HEPES buffered DMEM with 10% FBS. 24, 48, and 72 hours later, viral supernatant was collected and stored at 4°C and was replaced with fresh media after the first two collections. After the third collection, the viral supernatant was filtered with a 0.45 μM filter and was mixed with 1 volume Lenti-X concentrator to 3 volumes viral supernatant and incubated at 4°C for 1-3 days. Next, the virus was pelleted by centrifugation at 1,500 x g for 45 minutes at 4°C. The supernatant was removed, and the pellets were resuspended in HDF media to a final volume of 200 μL per 10 cm dish. The virus was stored for less than a week at 4°C until use. HDFs were seeded one day prior to transduction onto plates coated with 0.1% gelatin at the appropriate seeding density (e.g. 2.5k, 5k, or 10k per 96-well). HDFs were transduced two days in a row with 5 μL of each concentrated virus per 96-well. Fresh HDF media was included to reach a final volume of 100 μL per 96-well, and 5 ug/mL polybrene was added to increase transduction efficiency. Each day after the addition of virus, the plates were centrifuged at 1500 x g for 90 minutes at 32°C to further increase transduction efficiency via spinfection. At 1 dpi and 4-5 dpi, the media was replaced with fresh HDF media. At 7 dpi, the cells were dissociated using trypsin. Once the cells were in suspension, fresh HDF media was added, and the contents of each well were transferred to a laminin coated plate. The day after replating, the media was replaced with N3 media with RepSox. N3 media was replaced every 3 to 4 days until 35 dpi.

### Single cell RNA-sequencing

For scRNA-seq libraries generated for this work, cells were reprogrammed as described. Prior to loading onto 10X, cells were sorted via FACS with 1:1000 Zombie NIR™ spiked in prior to flow to sort for live cells. For the 4 dpi sample, a CellTrace-Far Red™ stain was also included starting at 1 dpi to sort for hyperproliferative cells. For 14 dpi samples, reprogrammed iMNs were also sorted via FACS for Hb9::GFP positive cells. Prior to loading, cells were resuspended, counted, confirmed for >90% viability by trypan blue staining, and loaded for a target recovery of ∼1,500 cells per condition. Libraries were generated using a 10X Chromium Next GEM Single Cell 3’ kit v3.1 (dual index). RT-PCR and Advanced Analytical Technologies Inc fragment analyzers were used to confirm pooled library quality. NovaSeq S4 kit was used to sequence pooled libraries with 150 reads for the cell barcode + UMI, 8 cycles for the i7 and i5 sample indices, and 150 cycles for the insert. Sequenced libraries were aligned to a custom reference based on GRCm39 with the eGFP marker gene added using CellRanger. ∼30-60k reads/cell were generated with the exception of LNI + DDRR reprogrammed cells at 4 dpi which had ∼5k reads/cell.

### Single cell RNA-seq analysis

#### Cluster analysis via Seurat

Cells were filtered for < 8% mitochondrial reads, and between 200 and 10,000 features. The SCTransform function was used to normalize libraries. An elbow plot was used to determine the number of PCA’s to use for clustering. Seurat’s FindNeighbors and FindClusters functions were used to cluster cells and the resolution was adjusted such that clusters largely belonged to reprogramming conditions. EmbMNs collected at E12.5 were clustered independently to identify median motor column (MMC) motor neurons. An MMC cluster was isolated using marker genes, Mnx1, Isl1, Lhx3, and eGFP (Davis-Dusenbery et al., 2014; Delile et al., 2019; Ichida et al., 2018; Stifani, 2014). For simplicity of notation, we denoted these simply “EmbMN” and ignored all non-MMC embMNs for analysis. After incomplete motor neurons were identified, this cluster was removed for further analysis.

### Surgery procedures for *in vivo* cell grafts

#### Surgery Procedures

All procedures were performed on mice under general anesthesia achieved through inhalation of isoflurane (1.5-3%) in oxygen-enriched air that was administered via a mouse anesthesia mask that was attached to a stereotactic device. Prior to surgery the head of the mouse was shaved and sterilely prepared with betadine and alcohol scrubs. All surgery procedures were conducted with the mouse stably positioned within a stereotactic frame using a mouse snout clamp, tooth bar, and dual sided ear bars. A heating pad positioned under the mouse was used to ensure maintenance of body temperature.

#### Craniotomy procedure

A single midline skin incision made using a scalpel was used to expose the dorsal surface of mouse skull and the surface of the skull is cleaned dry with a sterile cotton swab. Using a high-speed surgical drill, a small craniotomy was made over the left coronal suture that was visually aided by a stereomicroscope. As small rectangular flap of bone encompassing sections of the frontal and parietal bone was removed to expose the brain in preparation for injection. Saline was applied to keep the exposed brain tissue moisten throughout injection procedures.

#### Stroke lesion model

To create small focal ischemic strokes, 0.5 μL of L-NIO (N5-(1-Iminoethyl)-L-ornithine) (Cat. No. 400600-20MG, Millipore Sigma) (27.4 mg/ml (130 µM) in sterile PBS), an inhibitor of endothelial nitric oxide synthase, was injected into the caudate putamen nucleus at 0.2 μL/min using target coordinates relative to Bregma: +1.0 mm A/P, +2.5 mm L/M and −3.0 mm D/V using pulled borosilicate glass micropipettes (WPI, Sarasota, FL, #1B100-4). Glass micropipettes were mounted to the stereotaxic frame via specialized connectors and attached, via high-pressure polyetheretherketone (PEEK) tubing, to a 10 μL syringe (Hamilton, Reno, NV, #801 RN) controlled by an automated syringe pump (Pump 11 Elite, Harvard Apparatus, Holliston, MA). After the full volume of L-NIO solution had been injected, the pipette was allowed to rest at the injection site for 4 minutes before being slowly removed from the brain over the course of 1 minute. Following L-NIO injection, the surgical incision site was sutured and the animals were allowed to recover for 48 hours before undergoing iMN graft injections via a second surgery as described below.

#### iMN graft injections

iMN grafts were made in healthy mouse brain (n=5) and L-NIO stroke injured brain (n=5) at 48 hours after L-NIO injection. Healthy mice received a craniotomy prior to cell injections whereas the already created cranial window in stroke injured mice was not modified prior to cell injection. Two million imN cells recovered by FACS for Hb9::GFP-positive cells were resuspended in 70µL of basal media (N3, neurotrophic factors, RepSox, and penicillin-streptomycin) to create a concentrated cell solution at 28.6K cells/µL. The concentrated cell solution was backloaded into a pulled glass micropipette prior to connecting to the stereotaxic frame. The one cell stock was injected into all animals with injections alternating between mice from the healthy and stroke groups. 1 μL of cell solution was injected into the same striatal regions as that used to create the stroke at 0.15 μL/min using target coordinates relative to Bregma: +1 mm A/P, +2.5 mm L/M. The pipette was lowered to −3.5 mm D/V for the start of the injection and moved up +0.5mm twice over the course of the injection to a final location of −2.5 mm D/V to improve deposition of cells in brain tissue. The micropipette was allowed to dwell in the brain at the injection site for an additional 4 minutes at the end of the injection. The micropipette was then removed from the brain slowly and incrementally over a 1-minute period.

### Immunohistochemistry for *in vivo* cell grafts

#### Transcardial perfusions

After terminal anesthesia by overdose isoflurane inhalation, mice were perfused transcardially with heparinized saline and 4% paraformaldehyde (PFA) that was prepared from 32% PFA Aqueous Solution (Cat# 15714, EMS), using a peristaltic pump at a rate of 7 mL/min. Approximately, 10 mL of heparinized saline and 50 mL of 4% PFA was used per animal. Brains were immediately dissected after perfusion and post-fixed in 4% PFA for 6 hours at room temperature. After PFA post-fixing, brains were cryoprotected in 30% sucrose in Tris-Buffered Saline (TBS) at 4°C for at least 3 days with the sucrose solution replaced once after 2 days and stored at 4°C until further use.

#### Immunohistochemistry

Coronal sections of mouse brains receiving grafts were cut (40 µm thick) using a Leica CM1950 Cryostat. Tissue sections were stored in TBS buffer at 4°C in individual wells of a 96 well, plate prior to staining. Tissue sections were processed for immunofluorescence using free floating staining protocols described in comprehensive detail previously using donkey serum to block and triton X-100 to permeabilize tissue (O’Shea et al 2022 and Adewumi et al 2023). The primary antibodies used in this study were: rabbit anti-Hemagglutinin (HA) (1:1000, Sigma #H6908, RRID:AB_260070); goat anti-HA (1:800, Novus, NB600-362, RRID:AB_10124937); rat anti-Gfap (1:1000, Thermofisher, #13-0300, RRID:AB_86543); guinea pig anti-NeuN (1:1000, Synaptic Systems, 266 004, RRID:AB_2619988); rabbit anti-Neurofilament M (145kDa) (NFM) (1:500, Millipore Sigma, AB1987, RRID:AB_91201), chicken anti-GFP (1:2000, Abcam, ab13970, RRID:AB_300798), rabbit anti-GFP (1:500, Abcam, ab290, RRID:AB_303395), goat anti-Choline Acetyltransferase (ChAT) (1:500, MilliporeSigma, AB144P, RRID:AB_2079751) and rabbit anti-DARPP-32 (19A3) (1:400, Cell Signaling, 2306, RRID:AB_823479). All secondary antibodies used in this study were purchased from Jackson ImmunoResearch (West Grove, PA). All secondary antibodies were affinity purified whole IgG(H+L) antibodies with donkey host and target species dictated by the specific primary antibody used. All secondary antibodies were stored in 50% glycerol solution and diluted at a concentration of 1:250 in 5% normal donkey serum in TBS when incubated with brain histological sections. Nuclei were stained with 4’,6’-diamidino-2-phenylindole dihydrochloride (DAPI; 2ng/ml; Molecular Probes). Sections were mounted using ProLong Gold anti-fade reagent (ThermoFisher). Sections were examined and imaged using epifluorescence and deconvolution epifluorescence microscopy on an Olympus IX83 microscope. Orthogonal (3D) images were prepared using Zen 3.1 (Blue Edition) (Zeiss).

#### Quantification of immunohistochemistry

Quantification of immunohistochemistry staining intensity quantification was performed on whole brain section images taken with the 10X objective on the Olympus IX83 microscope. All images used for each comparable analysis were taken at a standardized exposure time and used the raw/uncorrected intensity setting. Quantification of antibody staining intensity was generated in the form of radial intensity profiles derived using NIH Image J (1.51) software and the radial profile angle plugin as detailed previously (O’Shea et al 2022 and Adewumi et al 2023). Total values for IHC stainings were determined by taking the integral (area under the curve) of the radial intensity profile.

### Cell coverslips

iMNs purified via FACS for Hb9::eGFP positive cells were cultured on 10 mm round coverslips (Thorlabs) coated with 0.1% gelatin in individual wells of a 24 well plate for 7 days following an initial seeding of 100,000 cells per well. iMNs were fixed with 4%PFA for 30 minutes and coverslips were then stained using IHC methods and antibodies described above.

## QUANTIFICATION AND STATISTICAL ANALYSIS

Quantification and statistical analysis were performed using Python and R. Code available upon request.

## FLUORESCENT IMAGING

Images not taken for immunohistochemistry were imaged using a Keyence™ All-in-one fluorescence microscope BZ-X800.

## ACKNOWLEDGEMENTS

Research reported in this manuscript was supported by the National Institute of General Medical Sciences of the National Institutes of Health under award number R35-GM143033. N.B.W. is supported by the National Science Foundation Graduate Research Fellowship Program under grant No. 1745302. The authors acknowledge the MIT SuperCloud and Lincoln Laboratory Supercomputing Center for providing HPC resources that have contributed to the research results reported within this paper. We thank the Koch Institute’s Robert A. Swanson (1969) Biotechnology Center (National Cancer Institute Grant P30-CA14051) for technical support, specifically the Flow Cytometry Core Facility. We thank Doug Lauffenberger and Christopher Johnstone, Sneha Kabaria, Emma Peterman, Deon Ploessl, Joji Teves, and Kasey Love for feedback on the development of the manuscript.

## AUTHOR CONTRIBUTIONS

N.B.W. and B.A.L. performed and analyzed murine reprogramming. B.A.L., A.M.B., and P.H. performed and analyzed human reprogramming. N.B.W. prepared and analyzed scRNA-seq libraries. N.B.W. and A.M.B. prepared iMNs for cell grafts. H.O.A. performed surgeries, staining, and analysis for cell grafts.

K.E.G. and T.M.O. supervised the project. N.B.W., K.E.G., B.A.L., H.O.A., and T.M.O. wrote the manuscript.

