## Supplementary Figures for "Proliferation history and transcription factor levels drive direct conversion"

**Fig. S1 Tailoring the conversion transcription factor module supports robust reprogramming**

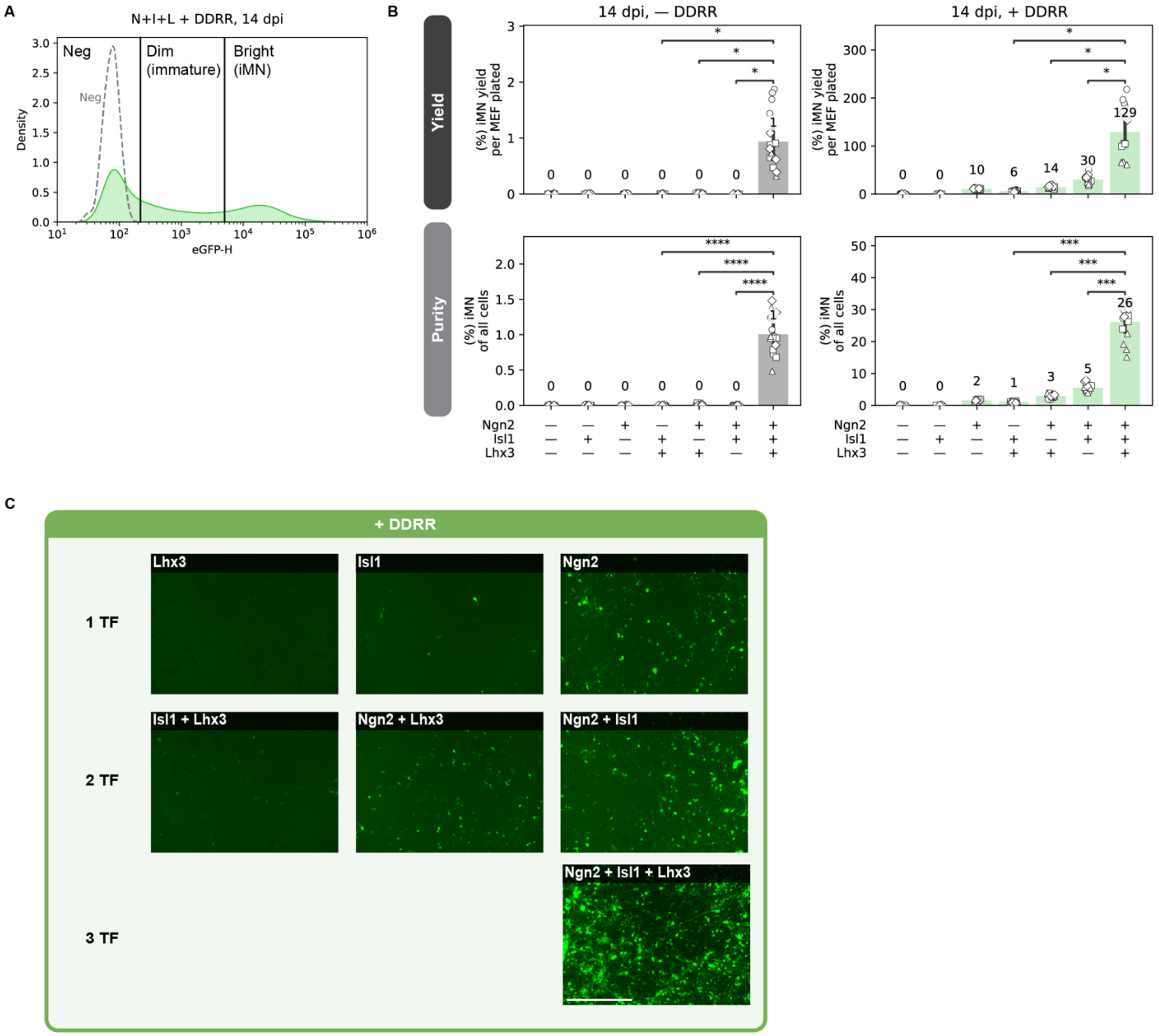


A. Histogram indicating the gate for quantifying iMNs based on Hb9::GFP fluorescence intensity using flow cytometry. Bright Hb9::GFP represent re iMNs (Babos et al., 2019) whereas dim cells represent immature or incomplete conversion states. Negative control are cells infected with just Lhx3.

B. Reprogramming purity and yield at 14 dpi for the transcription factor dropout experiment with Ngn2, Isl1, and Lhx3 ± DDRR. Mean is shown with 95% confidence interval; marker styles denotes biological reps; n = 4 biological reps per condition; one-tailed t-test with Bonferroni correction.

C. Images of the reprogrammed cells at 14 dpi for the transcription factor dropout experiment with Ngn2, Isl1, and Lhx3 ± DDRR. Scale bar represents 500 µm.

Significance summary: $p > 0.05$ (ns); *$p \leq 0.05$; **$p \leq 0.01$; ***$p \leq 0.001$; and ****$p \leq 0.0001$

**Fig. S2 Proliferation provides a principal axis to distinguish transcription factors' influence**

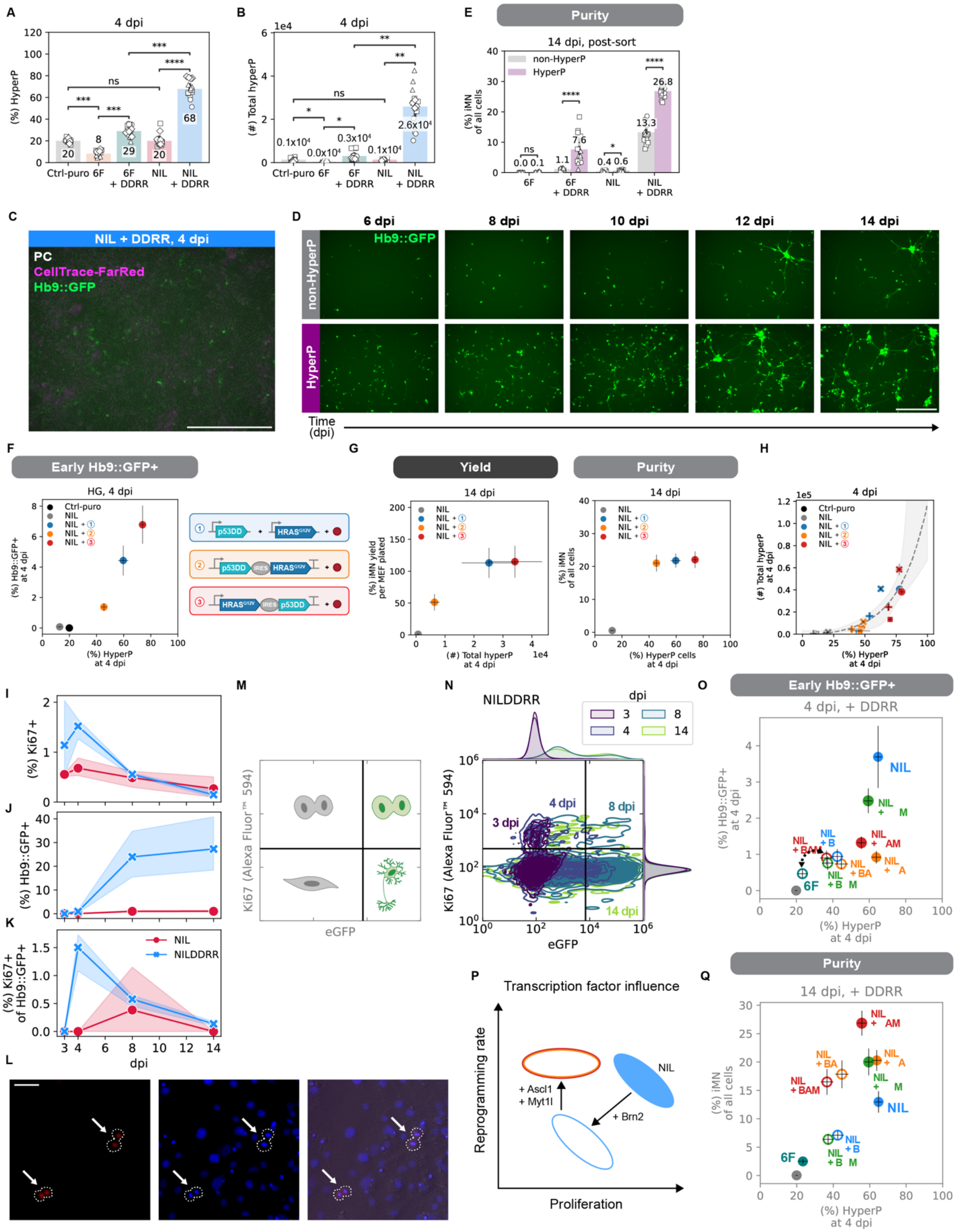

A-B. Hyperproliferative (HyperP) percent (A) and total number (B) at 4 dpi for 6F vs. NIL ± DDRR infected cells seeded at 10k/96-well. Ctrl-puro is 20% hyperP for a given biological rep by definition. Mean is shown with 95% confidence interval; marker styles denotes biological reps; n = 4 biological reps per condition; one-tailed t-test with Bonferroni correction.

C-D. Images of the labeled cells (see Fig. 2A for labeling strategy) for NIL + DDRR reprogrammed cells before sorting at 4 dpi (C) and after until 14 dpi (D). Both scale bars represent 500 μm.

A. Reprogramming purity at 14 dpi for cells sorted at 4 dpi into HyperP vs. non-HyperP populations within each condition. Mean is shown with 95% confidence interval; marker styles denotes biological reps; n = 3 biological reps per condition; one-tailed t-test.

B. Early Hb9::GFP reporter expression at 4 dpi vs hyperP percent at 4 dpi for different designs of the genetic portion of the DDRR cocktail. Mean is shown ± standard error of mean (SEM); n = 4 biological reps per condition.

G. Reprogramming purity at 14 dpi vs. hyperP percent at 4 dpi (left) and reprogramming yield at 14 dpi vs. hyperP total number at 4 dpi (right) for different designs of the genetic portion of the DDRR cocktail. Mean is shown ± standard error of mean (SEM); n = 4 biological reps per condition.

H. HyperP total number vs. percent at 4 dpi for different designs of the genetic portion of the DDRR. An exponential fit is shown with 95% confidence interval.

I-K. Ki67 staining from 3-14 dpi for NIL ± DDRR reprogrammed cells. The mean percent of cells that are Ki67+ (I), the percent of Hb9::GFP+ cells (J), and the percent of Hb9::GFP+ cells that are Ki67+ (K) are shown with 95% confidence interval.

L. Images of Ki67 staining at 4 dpi. Dashed lines demarcate Ki67+ cells that are going through mitosis/cytokinesis. Scale bar represents 50 μm.

M-N. Diagram depicting cell state categories (M) and representative contour plots (N) of Ki67 staining vs. eGFP fluorescence from 3-14 dpi for cells reprogrammed with NIL + DDRR.

O. Early Hb9::GFP reporter expression at 4 dpi in hyperP vs. non-hyperP populations within each reprogramming cocktail (all reprogrammed with +DDRR). Mean is shown with 95% confidence interval; marker styles denotes biological reps; n = 4 biological reps per condition; one-tailed t-test.

P. Diagram showing decrease in reprogramming and proliferation upon Brn2 addition. Addition of Ascl1 and Myt1l to Brn2-containing cocktails increases reprogramming yield but not proliferation.

Q. Reprogramming purity at 14 dpi vs. toal number of hyperP cells at 4 dpi for each reprogramming cocktail (all reprogrammed with +DDRR). Mean is shown ± standard error of mean (SEM); n = 4 biological reps per condition. Black dashed line connects 6F when NIL is encoded on 3 separate viruses (i.e. B+A+M+N+I+L) vs. when NIL is on a single polycistronic transcript (i.e. B+A+M+NIL).

Significance summary: $p > 0.05$ (ns); *$p \leq 0.05$; **$p \leq 0.01$; ***$p \leq 0.001$; and ****$p \leq 0.0001$

**Fig. S3 Titration of individual transcription factors reveals factor-specific influence on conversion rates**

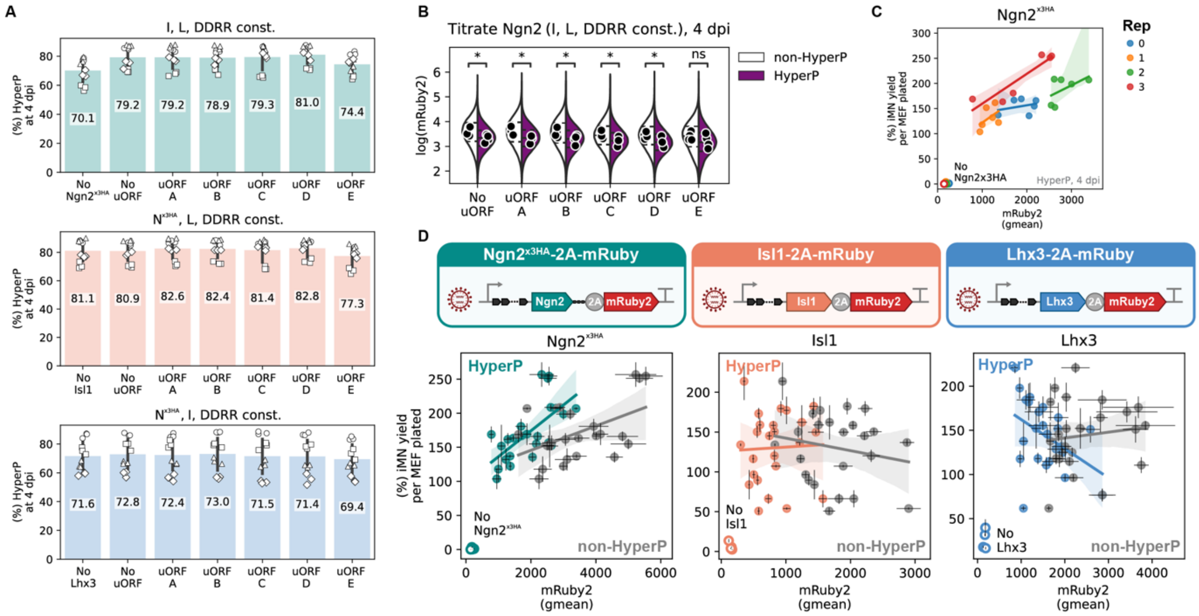


A. Percent hyperP cells across uORF conditions for each titrated TF at 4 dpi. Mean is shown with 95% confidence interval; marker styles denote biological reps; n = 4 biological reps per condition.

B. A representative plot using Ngn2$^{x3HA}$ showing how mRuby2 expression at 4 dpi is lower in hyperP than non-hyperP cells across uORFs. Geometric mean is shown by markers and distribution is shown by violin plot with inter-quartile range marked by dashed lines; n = 4 biological reps per condition; one-tailed t-test.

C. Same reprogramming yield data vs. mRuby2 expression in 4 dpi hyperP cells as Fig. 3D but split by biological rep to show how heterogeneity in transgene expression across biological reps can be leveraged to explore larger ranges of expression.

D. Same reprogramming yield data vs. mRuby2 expression in 4 dpi cells but split by non-hyperP (grey) vs. hyperP (colored) cells to show how proliferation affects reprogramming correlations.

Significance summary: p > 0.05 (ns); *p ≤ 0.05; **p ≤ 0.01; ***p ≤ 0.001; and ****p ≤ 0.0001

**Fig. S4 Expression of reprogramming factors varies across polycistronic cassettes**

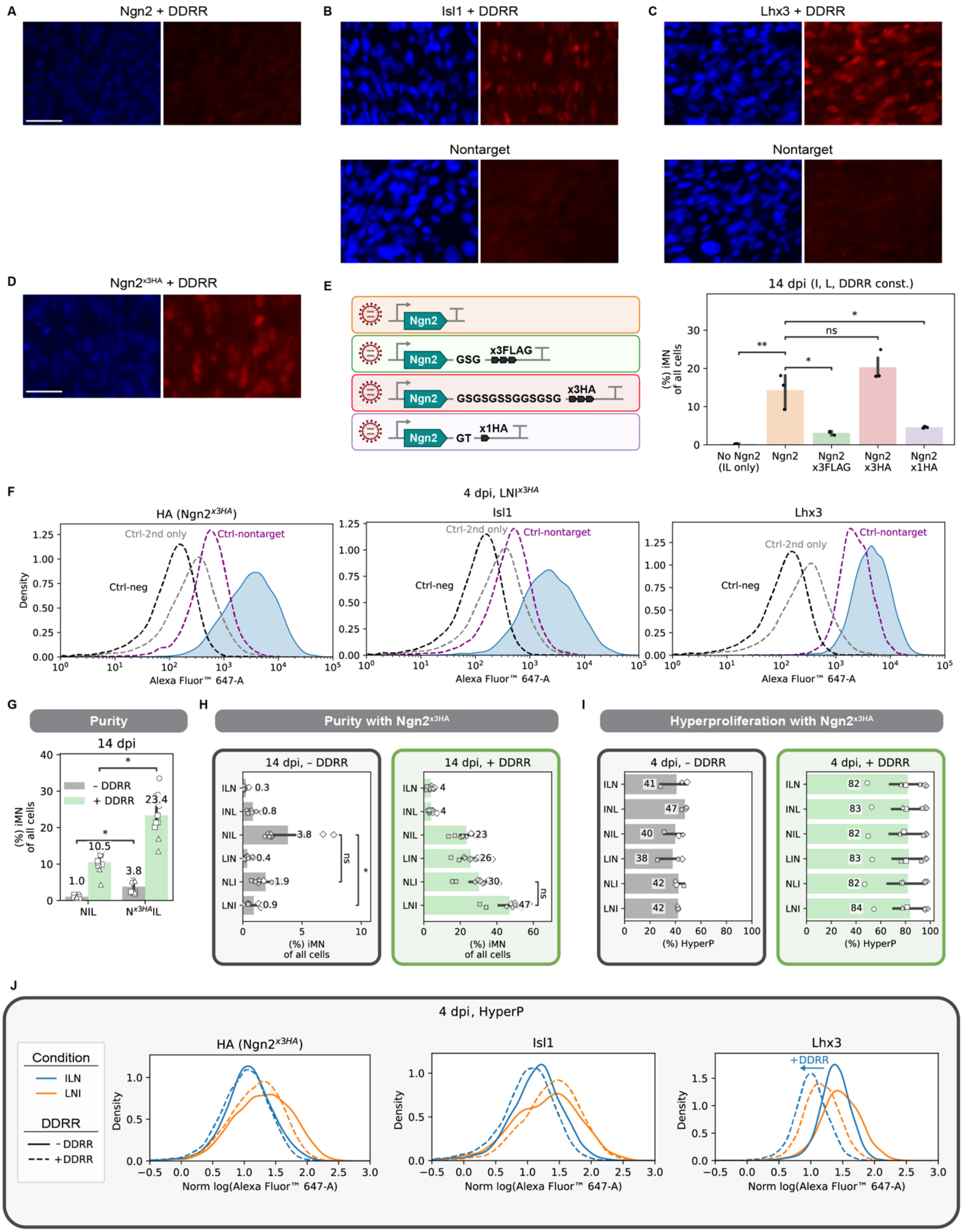

A-C. Images demonstrating Isl1 and Lhx3 antibodies properly localize to specific to expression of the transgene while Ngn2 fails. Nontargeting control for Isl1 antibody was Lhx3 + DDRR and nontargeting antibody for Lhx3 was Isl1 + DDRR. Scale bar represents 50 μm.

D. Image demonstrating HA staining properly localizes to the nucleus for cells infected with $Ngn2^{x3HA}$. Scale bar represents 50 μm.

E. Reprogramming yield at 14 dpi for Ngn2 with different linker-tags. Mean is shown with 95% confidence interval; two-tailed t-test with Bonferroni correction.

F. Histograms of immunofluorescence from staining for HA ($Ngn2^{x3HA}$), Isl1, and Lhx3 measured via flow cytometry. Control negative, secondary stain only, and a nontargeting control (cells infected with just +DDRR) are shown in black, grey, and purple dashed lines, respectively.

G. Reprogramming purity at 14 dpi for the polycistronic NIL cassette ± DDRR for the untagged Ngn2 and the tagged $Ngn2^{x3HA}$.

H. Reprogramming purity at 14 dpi for the different polycistronic TF cassettes without DDRR (left) and with +DDRR (right). All cassettes contain the tagged $Ngn2^{x3HA}$. Mean is shown with 95% confidence interval; marker styles denotes biological reps; n = 3 biological reps per condition; one-tailed t-test with Bonferroni correction.

I. Percent hyperP cells for the different polycistronic transcription factor cassettes without DDRR (left) and with +DDRR (right) at 4 dpi. Mean is shown with 95% confidence interval; markers denote biological reps.

J. Histograms of immunofluorescence at 4 dpi for a couple representative polycistronic cassettes (ILN: blue, and LNI: orange) without DDRR (solid) and with +DDRR (dashed).

Significance summary: $p > 0.05$ (ns); *$p \leq 0.05$; **$p \leq 0.01$; ***$p \leq 0.001$; and ****$p \leq 0.0001$

**Fig. S5 Driving hyperproliferation improves direct conversion of adult human fibroblasts**

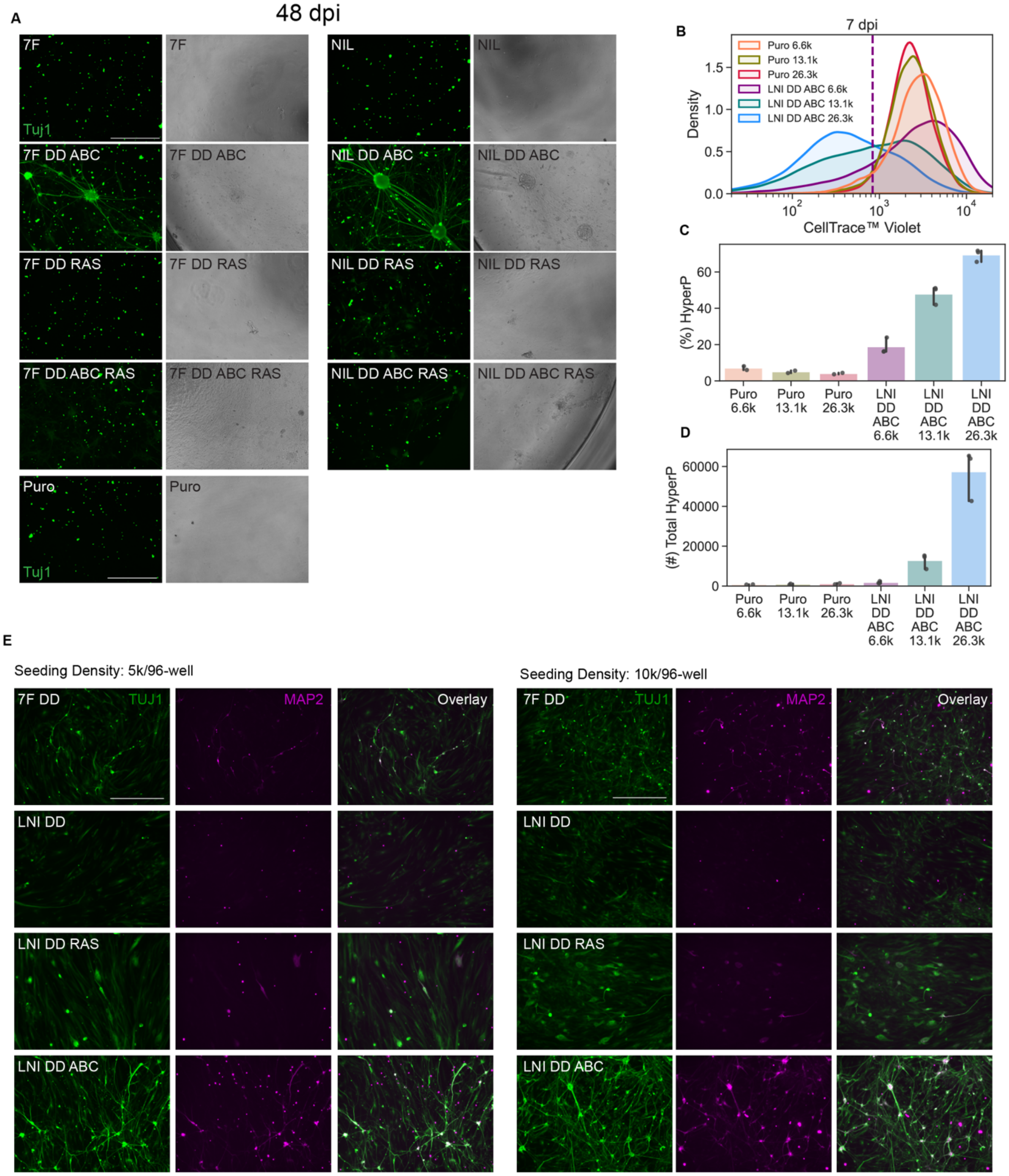


A. Representative images at 48 dpi of cells fixed and stained with TUJ1 and corresponding brightfield images. Scalebar represents 500 µm.
B. Representative CellTrace distributions for Puro and LNI DD ABC across different seeding densities per 48-well. Hyperproliferative cells are defined relative to the 5%-lowest fluorescent cells in all control puro seeding densities.
C. Hyperproliferative (HyperP) percent at 7 dpi across conditions. Ctrl-puro is 5% HyperP by definition. Mean is shown with 95% confidence interval; n=1 biological reps per condition; One-tailed t-test with Bonferroni correction.
D. Hyperproliferative (HyperP) total number at 7 dpi across conditions. Mean is shown with 95% confidence interval; n=1 biological reps per condition; One-tailed t-test with Bonferroni correction.
E. Representative images at 35 dpi of cells fixed and stained with TUJ1 and MAP2 for different seeding densities. Scalebar represents 500 µm.

**Fig. S6 Compact cocktail enables neurotrophic factor-free reprogramming**

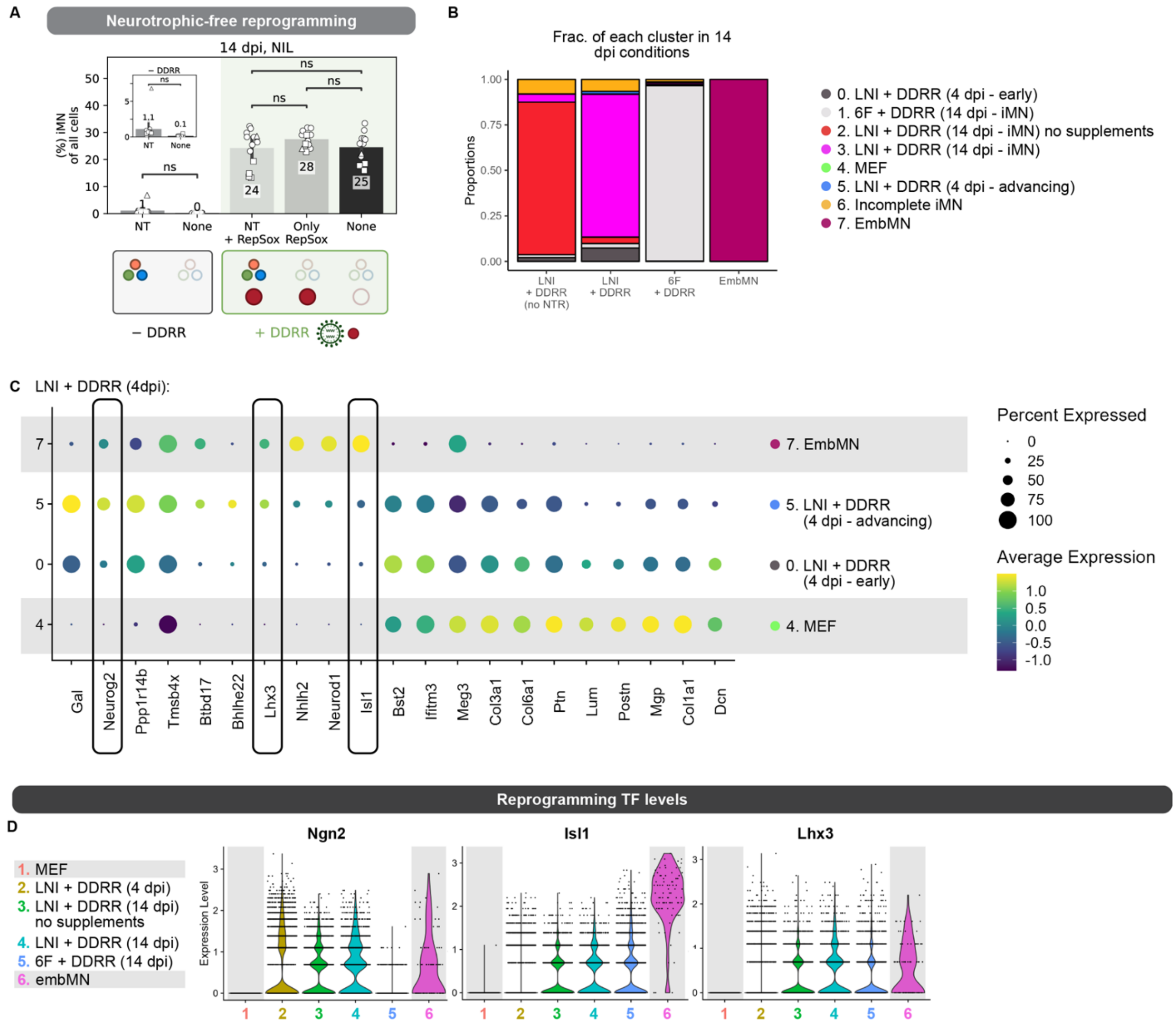


A. Reprogramming purity at 14 dpi for NIL ± DDRR. NIL infected cells were reprogrammed in N3 media ± neurotrophic growth factors (NT). NIL + DDRR infected cells were reprogrammed in N3 media with both NT and RepSox, with only RepSox, and with neither. Inset shows iMN purity for NIL (without DDRR) with zoom-in axis for clarity. Mean is shown with 95% confidence interval; marker styles denotes biological reps; n = 3 biological reps per condition; One-tailed t-test with Bonferroni correction.

B. Distribution of embMNs and iMNs at 14 dpi across the 8 identified clusters (0-7) based on (Fig. 5D). Each library primarily maps to a single cluster.

C. Feature plot for the 21 genes (top 20 marker genes and Isl1) differentiating the two major clusters in 4 dpi hyperP cells reprogrammed with LNI + DDRR. MEFs and embMN (clusters 4 and 7, respectively) are included as baselines. The iMN reprogramming factors (Ngn2, Isl1, Lhx3) are outlined with a box.

D. Violin plots of expression across reprogramming conditions for the reprogramming TFs. Cells grouped in the incomplete iMN (cluster 6) are removed from 14 dpi iMNs to allow comparison of fully converted cells.

Significance summary: p > 0.05 (ns); *p ≤ 0.05; **p ≤ 0.01; ***p ≤ 0.001; and ****p ≤ 0.0001

**Fig. S7 Compact cocktail generates engraftable neurons**

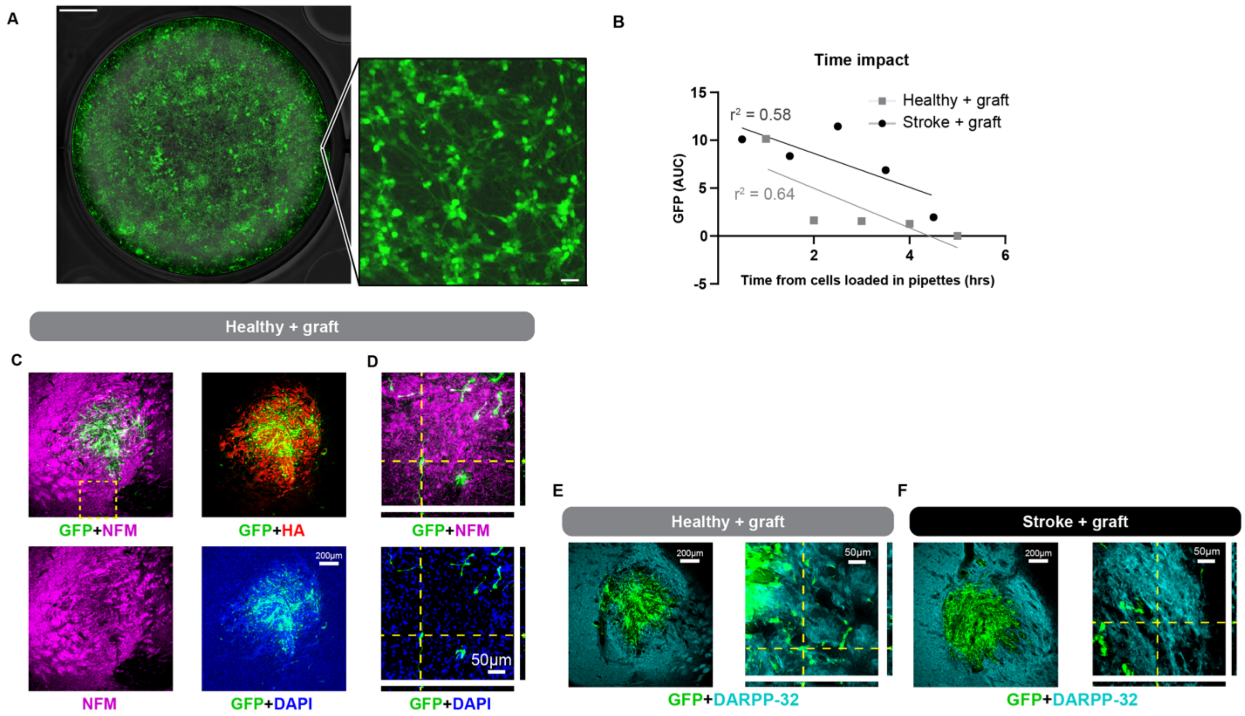


A. Image of reprogrammed iMNs in a 6-well at 14 dpi demonstrating that reprogramming induced with LNI + DDRR can scale with well-size. Phase contrast and Hb9::GFP overlaid and an inset confirms neuronal morphology. Scale bar represents 5 mm in larger image and 50 μm in the smaller inset.

B. Correlation analysis of total GFP at 350 microns vs. time of graft relative to when cell grafts were loaded in pipettes with $r^2$ values.

C-D. Coronal sections of healthy mouse striatum 2 weeks after reprogrammed iMN cell engraftment at various objectives. Sections were imaged for GFP, neurofilament (NFM), and NeuN.

E-F. Coronal sections of healthy (E) and stroke-induced (F) with grafted iMNs stained with striatal neuron marker (DARPP-32).
